# *JUN* upregulation drives aberrant transposable element mobilization, associated innate immune response, and impaired neurogenesis in Alzheimer’s disease

**DOI:** 10.1101/2022.11.24.517794

**Authors:** Chiara Scopa, Samantha M. Barnada, Maria E. Cicardi, Mo Singer, Davide Trotti, Marco Trizzino

## Abstract

Adult neurogenic decline, inflammation, and neurodegeneration are phenotypic hallmarks of Alzheimer’s disease (AD). Mobilization of transposable elements (TEs) in heterochromatic regions was recently reported in AD, but the underlying mechanisms are still underappreciated. Combining functional genomics with differentiation of familial and sporadic AD patient derived-iPSCs into hippocampal progenitors, CA3 neurons, and cerebral organoids, we found that upregulation of the AP-1 subunit c-JUN triggers decondensation of genomic regions containing TEs. This leads to cytoplasmic accumulation of TE-derived RNA-DNA hybrids, activation of the cGAS-STING cascade, and increased cleaved caspase-3 levels, suggesting initiation of programmed cell death in progenitor cells and neurons. Notably, inhibiting c-JUN effectively blocks all the downstream molecular processes and rescues neuronal death and impaired neurogenesis in the AD progenitors. Our findings open new avenues for identifying therapeutic strategies and biomarkers to counteract disease progression and diagnose AD in the early, pre-symptomatic stages.

## Introduction

Alzheimer’s disease (AD), the most frequent cause of dementia^1–3^, is an age-related neurodegenerative disorder characterized by progressive memory loss and decline of cognitive functions. AD’s histopathological hallmarks include extracellular Amyloid beta (Aβ) plaques and intracellular neurofibrillary TAU tangles (NFTs)^4^. One of the first brain regions to show these pathological features is the hippocampus^5^. The subgranular zone of the dentate gyrus within the hippocampus is a human neurogenic niche^6–8^ which harbors neural stem cells and controls cell fate determination^9^. Interestingly, defects in adult hippocampal neurogenesis are associated with various neurodegenerative disorders, including AD^10–13^.

Several studies report alterations in hippocampal neurogenesis in transgenic animal models of AD^12^ and an exacerbated decline of adult neurogenesis in AD patients^14,15^. Notably, these alterations occur in the early stages of the disease^12^, suggesting they play a role in triggering the onset of the disease’s clinical phenotypes. Impaired neurogenesis likely accelerates and facilitates neurodegenerative progression^16^. Moreover, these studies highlight the link between the key molecules of AD (i.e. TAU and Aβ) and neurogenesis. TAU is involved in the microtubule dynamics required for axonal outgrowth and plays an essential role in hippocampal neurogenesis^17^. Recent studies indicate that TAU hyperphosphorylation impairs hippocampal neurogenesis^18,19^.

Increasing evidence suggests that intracellular accumulation of Aβ also negatively impacts neural precursors cell (NPC) proliferation and neuronal differentiation during hippocampal neurogenesis^20–22^. Additionally, neural stem cell (NSC) fate determination, and therefore neurogenesis, is regulated by MAP kinases^23–26^. Interestingly, several studies suggest a compelling link between MAPK signaling and AD pathogenesis, revealing that the c-JUN-amino-terminal kinase (JNK) pathway is involved in Aβ-induced neurodegeneration^27–30^ and in the hyperphosphorylation of TAU contributing to the formation of the NFTs^31–33^. Importantly, c-JUN is the downstream effector of the JNK pathway.

Phospho-c-JUN is a fundamental member of the AP-1 family of transcription factors, functioning as either homodimers (c-JUN/c-JUN) or as heterodimers (c-JUN/c-FOS, c-JUN/ATF2, c-JUN/MAF)^34^. Among various other functions, AP-1 modulates cell death signals^31,32^ and promotes the transcription of a series of pro-apoptotic factors, such as TNF-α, FAS-L, c-MYC and ATF3, which induce cell death by apoptosis^31–33^. Recent lines of evidence suggests that AP-1 can act as a pioneer factor by binding condensed nucleosomes and recruiting the BAF complex to elicit chromatin accessibility^35–37^. Notably, upregulation of *JUN* (which encodes for c-JUN) has been detected in many neurodegenerative diseases, including AD^32,38^. Nonetheless, the link between aberrant c-JUN activity and the associated neurodegenerative outcomes have not been explored in depth.

Finally, there is mounting evidence indicating a role for transposable elements (TEs) in the molecular pathogenesis of AD. More specifically, the disease is characterized by aberrant de-repression and mobilization of TEs found in regions of repressed chromatin, particularly retrotransposons belonging to the long interspersed nuclear elements (LINEs) and long terminal repeat containing transposons (LTRs)^39–46^. Yet, the mechanisms leading to TE de-repression and the functional consequences of this phenomenon in AD pathogenesis are understudied, especially in humans.

In this study, we leveraged familial and sporadic AD patient-derived induced pluripotent stem cells (iPSCs) and differentiated them into hippocampal progenitors, CA3 neurons and cerebral organoids. We demonstrated that c-JUN is the upstream regulator of the transcriptional network altered in AD hippocampal progenitors, and that the aberrant upregulation of *JUN* leads to the de-repression and mobilization of hundreds of TEs. Moreover, we found that aberrant TE mobilization induces a cytoplasmic accumulation of RNA-DNA hybrids, which elicits the activation of the cGAS-STING pathway and, ultimately, activation of caspase-3, suggesting the initiation of programmed cell death. Inhibiting c-JUN phosphorylation blocks this cell-death axis in AD progenitors by maintaining TE repression, ultimately preventing the activation of the downstream pathogenic cascade.

## Results

### iPSC-derived model of human neurogenesis in familial Alzheimer’s disease

To investigate the role of c-JUN in the onset of Alzheimer’s disease (AD), we started from familial AD and then replicated our findings in sporadic AD. Specifically, we derived hippocampal precursor neurons from AD patients and control human iPSCs. First, we confirmed the pluripotent state and the expression of *JUN* (which encodes the c-JUN protein) in two control (CTRL1 and CTRL2) and two familial AD lines (FAD1 and FAD2). CTRL and FAD lines included both sexes and were derived from individuals of comparable age. The FAD1 cell line contains an *APP* gene duplication and the FAD2 line contains a heterozygous missense mutation in *PSEN2* (*PSEN2*: p.Asn141Ile).

The immunofluorescence analysis for the pluripotency markers, NANOG and OCT4, and the immunofluorescence for c-JUN showed that the fraction of cells expressing these markers was not significantly different between CTRL and AD lines, indicating that the AD lines are pluripotent and they have the same percentage of JUN expressing cells than the control lines (**Extended Data Fig. 1a, b**). However, FAD iPSCs showed higher expression levels of c-JUN (**Extended Data Fig. 1b)**.

We then differentiated the four iPSC lines (CTRL1, CTRL2, FAD1, and FAD2) to hippocampal neural precursor cells (hpNPCs) using a previously published protocol (**Fig. 1a**)^47^. To assess the composition of the obtained hpNPC population, we quantified the expression of the established markers for different hippocampal neural precursor stages via qPCR (Fig. 1b). NESTIN defines early precursors, TBR2 and FOXG1 define intermediate progenitors, PROX1 defines late progenitors, and DCX defines neuroblasts^48^ (**Fig. 1b**). We observed that the FAD hpNPCs have impaired neurogenesis, indicated by a robust reduction of early neural stem cells (NSCs; NESTIN-positive cells) and a substantial increase in TBR2- and FOXG1-positive intermediate progenitors (**Fig. 1b**). This result was also confirmed by immunofluorescence (**Fig. 1c, d**). Moreover, the FAD hpNPCs failed to properly differentiate into neuroblasts (**Fig. 1b, d**), demonstrated by the significantly reduced expression of the neuroblast marker, DCX, as well as the reduced percentage of the DCX positive cells (**Fig. 1c**).

**Figure 1.**
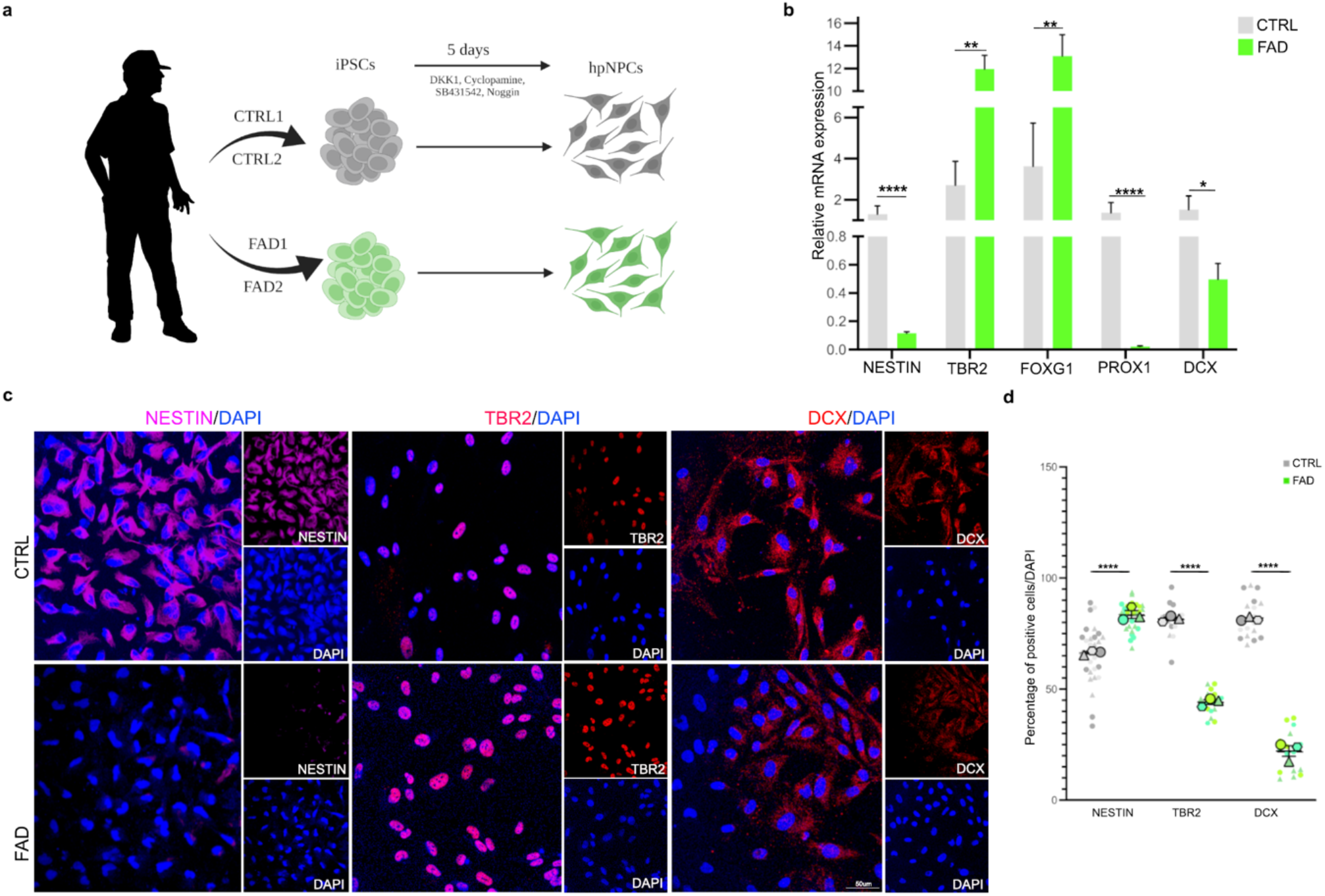
FAD iPSC-derived hippocampal neural progenitors display impaired neurogenesis. **a**. Scheme of the protocol for hippocampal neural progenitor cell (hpNPC) differentiation (made with BioRender.com). iPSCs were derived from the skin fibroblasts of two patients with familial Alzheimer’s disease (FAD1 and FAD2) and two healthy controls (CTRL1 and CTRL2), and differentiated into hpNPCs after 5 days in induction media. The hpNPCs were maintained in proliferation media post-induction. **b, c**, & **d**. qPCR and immunofluorescence for markers of different stages of hpNPC populations. The qPCR shows a neurogenic defect in FAD progenitors, with enrichment for TBR2-positive intermediate progenitors as also confirmed in the immunofluorescence and relative quantification for hpNPC population markers. NESTIN – early precursors; TBR2/FOXG1 – intermediate progenitors; PROX1 – late progenitors; DCX – neuroblasts. Scale bar 50um, 40X magnification. DAPI staining on nuclei in blue.

### *JUN* regulates the transcriptional network dysregulated in familial Alzheimer’s disease hippocampal neural progenitors

To further investigate the differences between healthy and FAD hippocampal neural progenitors, we characterized their transcriptomes using RNA-sequencing (RNA-seq). After 20 days in proliferation medium, we collected the cells to perform RNA-seq. This analysis identified 1,976 differentially expressed genes, 751 of which (38.1%) were downregulated, and 1,225 (61.9%) were upregulated in the FAD progenitors (FDR <5%; log2(FC) ± 1.5; **Fig. 2a**). In agreement with RT-qPCR and immunofluorescence data (**Fig. 1**), the early progenitor markers, *NESTIN* and *PAX6*, were downregulated, confirming confirming the impaired neurogenesis in FAD progenitors.

**Figure 2.**
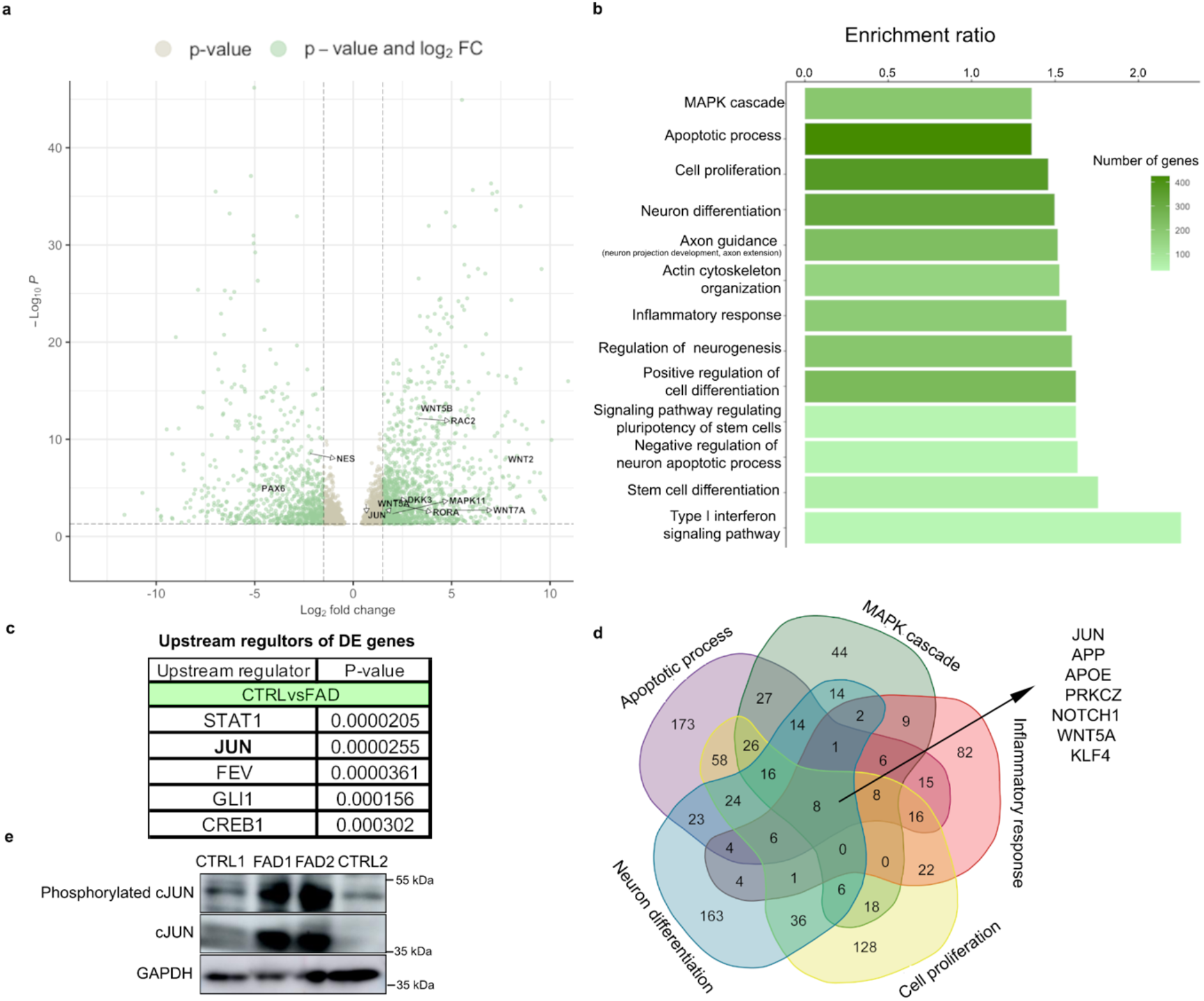
*JUN* upregulation causes dysregulated transcriptional networks in FAD hpNPCs. **a**. Volcano plot showing genes differentially expressed in FAD hpNPCs relative to CTRL hpNPCs. Labeled differentially expressed genes are involved in neurogenesis (*PAX6, NES*) and WNT/JNK signaling (*WNT2, WNT7A, WNT5B, RAC2, RORA, MAPK11, DKK3, WNT5A*). Green = differentially expressed genes passing significance thresholds p-value < 0.05 and log2(fold-change) +/– 1.5; Gray = differentially expressed genes passing significance threshold of p-value < 0.05. **b**. Enriched pathways associated with the 1,976 differentially expressed genes in FAD hpNPCs predicted by WebGestalt. **c**. Top upstream regulators/transcription factors of the 1,976 differentially expressed genes in FAD hpNPCs, as predicted by Ingenuity Pathway Analysis (Qiagen). **d**. Venn diagram showing the genes shared across all of the top five enriched pathways with the most differentially expressed genes (venn diagram made with https://bioinformatics.psb.ugent.be). **e**. Immunoblot displaying the upregulation of both c-JUN and phosphorylated c-JUN across FAD and CTRL hpNPCs.

Several studies have demonstrated a critical role for WNT signaling in the pathogenesis of AD^49–55^ and in the regulation of adult hippocampal neurogenesis^56–63^. Accordingly, the expression of several genes involved in both the canonical (*DKK3, WNT7A, SFRP4, WNT2*) and non-canonical (*WNT5A, RORA, RAC2*) WNT signaling pathways were upregulated in FAD hpNPCs (**Fig. 2a**). Moreover, *DKK1* was expressed more in the FAD lines relative to the CTRLs (**Extended Data Table 1**). DKK1 is an antagonist of the canonical WNT signaling pathway^64^, which leads to the activation of the WNT/JNK pathway, which ultimately results in increased phosphorylation of c-JUN (encoded by *JUN*)^65^. Furthermore, the expression of the *JUN* gene itself was significantly upregulated in the FAD lines (log2(FC) = -0.6611642; p-value = 0.00039024; **Fig. 2a**). Consistent with this, the activation of the WNT/JNK pathway was suggested to play a role in Aβ oligomer neurotoxicity^51,66^.

We employed the WEB-based GEne SeT AnaLysis Toolkit (WebGestalt)^67^ to identify pathways associated with the 1,976 differentially expressed genes. This analysis revealed that these genes are associated with inflammation, neurogenesis, and neural differentiation, as well as cytoskeleton organization, apoptotic process, and the MAPK cascade (**Fig. 2b**). Notably, Ingenuity Pathway Analysis (Qiagen) identified c-JUN as one of the top enriched transcriptional regulators upstream to all of the differentially expressed genes, suggesting that most of the differentially expressed genes are c-JUN targets (**Fig. 2c**). Moreover, when analyzing the top enriched pathways that included the greatest number of differentially expressed genes, *JUN* was one of only 8 genes shared across all these pathways (Fig. 2d). To functionally test this computational prediction, we performed an immunoblot confirming upregulation of c-JUN and phosphorylated c-JUN in FAD hpNPCs (**Fig. 2e**)

Overall, these results indicate that the FAD progenitors are characterized by aberrant activation of the WNT/JNK pathway, an upregulation of *JUN* expression (which encodes for c-JUN), and dysregulation of its target genes.

### The WNT/JNK pathway is dysregulated in FAD CA3 hippocampal neurons

Recent studies have demonstrated that canonical WNT signaling is inhibited by several pathogenic mechanisms in the brain of AD patients which leads to neural death and synaptic plasticity regulation impairment^53,54^. To investigate whether aberrant activation of the WNT/JNK pathway was also occurring in AD neurons, we differentiated CTRL and FAD progenitors into CA3 hippocampal neurons using an established protocol (**Extended Data Fig. 2a**)^68^. Both CTRL and FAD differentiated neurons expressed the specific CA3 markers, such as Glutamate Ionotropic Receptor Kainate Type Subunit 4 (GRIK4) and Secretagogin (SCGN; **Extended Data Fig. 2b**). However, the differentiation of the FAD lines resulted in a reduced number of mature CA3 neurons relative to the CTRL (51.4% in FAD; 87.2% in CTRL, **Extended Data Fig. 2b**).

We performed RNA-seq on the CA3 neurons and identified 1,105 differentially expressed genes between CTRL and FAD (FDR < 5%; **Extended Data Fig. 2c**). 82 of these genes, including *MAPK10* (*JNK3*), *ROR2, WNT7A* and *WNT7B*, are involved in both the WNT/JNK pathway and the MAPK cascade. Notably, using WebGestalt we identified the MAPK cascade as one of the top 10 differentially expressed pathways using WebGestalt (**Extended Data Fig 2d**).

Finally, when analyzing the top enriched pathways that included the greatest number of differentially expressed genes, we found that only *MAPT* is shared across these pathways (**Extended Data Fig. 2e**). *MAPT* encodes for TAU, one of the key proteins in AD pathogensis^69,70^, and recent studies have reported a correlation between aberrant c-JUN activity and the formation and maturation of NFTs^32^.

In summary, we observed that the WNT/JNK pathway is dysregulated, not only in FAD hippocampal progenitors but also in the CA3 hippocampal neurons CA3 neurons, suggesting a critical role for c-JUN in both these cell types.

### Dysregulated chromatin accessibility in FAD hippocampal progenitors

To investigate if the transcriptomic aberrations identified in the FAD lines are associated with significant differences in chromatin accessibility, we performed an ATAC-seq on the CTRL- and FAD-derived progenitors, generating 150 bp long Paired-End reads.

We identified 3,100 differentially accessible (DA) regions between CTRL and FAD hpNPCs (FDR <5%; log2(FC) ±1.5). Of these regions, 68.35% were accessible in FAD compared to CTRL (**Fig. 3a**: FAD Up). By examining the closest gene to each of the 3,100 DA regions, we found that approximately 14% (430) of these DA regions were located nearest to a differentially expressed gene (i.e. their nearest gene was differentially expressed; **Fig. 3b**), suggesting that there are at least 430 enhancer-gene pairs (or promoter-gene pairs) dysregulated in the FAD progenitors. Of the 430 DA regions, 94.1% were putative enhancers (distance from closest transcription start site or TSS > 1kb; **Fig. 3c**) whereas 5.9% were putative promoters (TSS distance < 1kb).

**Figure 3.**
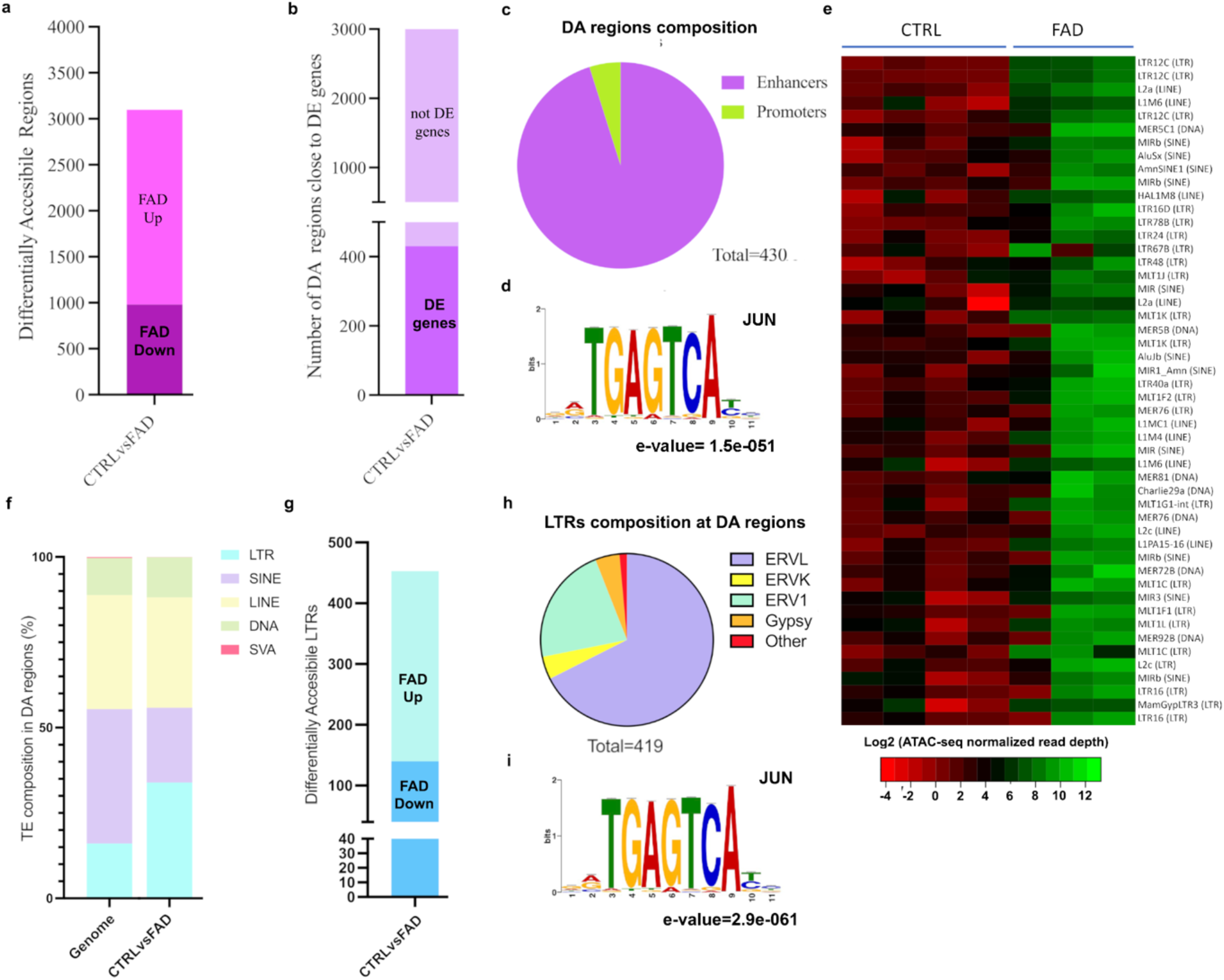
Differentially accessible transposable elements in FAD hpNPCs. **a**. Differentially accessible (DA) regions in the FAD hpNPCs compared to CTRL hpNPCs. FAD Up = significantly more accessible in FAD relative to CTRLs; FAD Down = significantly less accessible in FAD relative to CTRLs. **b**. DA regions located near a differentially expressed gene. **c**. DA regions near differentially expressed genes are predominantly enhancers (>1 kb from the transcription start site or TSS). **d**. MEME-ChIP analysis of 3,100 DA regions uncovered an enriched binding motif for JUN. **e**. Heatmap showing the top 50 TE copies identified as significantly more accessible in FAD relative to CTRL hpNPCs. **f, g, h**. Family distribution of the aberrantly active TEs in FAD progenitors. **i**. The aberrantly active LTRs are enriched for the JUN motif.

We then performed DNA-based motif analysis (MEME-ChIP) to identify any potential transcription factors underlying the changes in chromatin accessibility in the 3,100 DA regions. Remarkably, the most significantly enriched binding motif was the sequence recognized by JUN (e-value=10^−51^; **Fig. 3d**).

### Transposable elements are aberrantly active in FAD hippocampal progenitors

Aberrant de-repression of transposable elements is a hallmark of AD^42,46^. In agreement, we observed that 1,336 DA regions overlapped with a TE. The top 50 TE copies identified as significantly more accessible (i.e. active) in FAD relative to CTRL progenitors were predominantly (82%) retrotransposons (RTEs). Of these re-activated RTEs, 53.6% were LTRs, while the remaining were more or less equally distributed between LINEs (long interspersed nuclear elements) and SINEs (short interspersed nuclear elements; **Fig. 3e**). This aberrant LTR mobilization in the DA regions of the FAD lines was also investigated at the TE family level (**Fig. 3f**). In detail, 93.4% of the differentially accessible LTRs were endogenous retroviruses (ERVs; **Fig. 3h**). In line with these findings thus far, a motif analysis conducted on just the differentially accessible LTRs also revealed enrichment for the binding motif of JUN (e-value =10^−61^, **Fig. 3h**).

Together, these data indicate that the FAD progenitors are characterized by dysregulated chromatin accessibility at thousands of genomic sites, including hundreds of transposable elements. Moreover, our data suggests that most of these genomic regions are c-JUN target sites, and that aberrant c-JUN activity may underlie this observed chromatin dysregulation.

Several recent studies have demonstrated that AP-1 can act as a pioneer transcription factor by recruiting the BAF chromatin remodeling complex at condensed genomic regions to elicit chromatin accessibility^35,71–73^. Thus, we suggest that aberrantly active AP-1 in FAD progenitors may recruit BAF to its target sites (i.e. at the regions harboring the AP-1 motif), activating the TEs within these genomic regions.

### Aberrant TE mobilization elicits cGAS-STING activation

Several studies have demonstrated that aberrant TE de-repression and mobilization is observed across a variety of neurodegenerative disorders, including AD^39–41,43–46^. A recent study in blind mole rats (an ageing model for longevity studies) has shown that TE de-repression leads to the cytosolic accumulation of TE-derived RNA-DNA hybrids, which activates the cGAS-STING innate immune pathway leading to cell death^74^. The blind mole rats leverage this mechanism to suppress pre-cancerous cells^74^. We thus set out to investigate if the aberrant TE de-repression observed in the AD brains may also lead to cytosolic RNA-DNA hybrid accumulation, innate immune response and apoptosis, all of which are hallmarks of the disease.

Intriguingly, immunostaining conducted on FAD and CTRL progenitors with the S9.6 antibody, specific for the detection of RNA-DNA hybrids^74^, displayed significant hybrid accumulation in the cytoplasm of both FAD lines relative to the CTRLs (**Fig. 4a, b**). The RNA-DNA hybrid accumulation could be facilitated by the reduced expression levels of Ribonuclease H (RNase H) observed in the FAD progenitors. Namely, RNase H is specialized to selectively degrade RNA-DNA hybrids present in cells (**Fig.4c**)^75^. Additionally, we detected a robust increase in STING immunofluorescence signal quantified in the FAD progenitors (**Fig. 4a, b**). The upregulation of STING and cGAS in the FAD progenitors was further validated by western blot (**Fig. 4d**). Because activation of the cGAS-STING pathway in cells leads to their death via apoptosis^76^, we assessed whether FAD progenitors were executing this cell death program. Using western blot analysis, we observed an increase in cleaved caspase 3 (CC3) in the FAD lines compared to controls (**Fig. 4d**).

**Figure 4.**
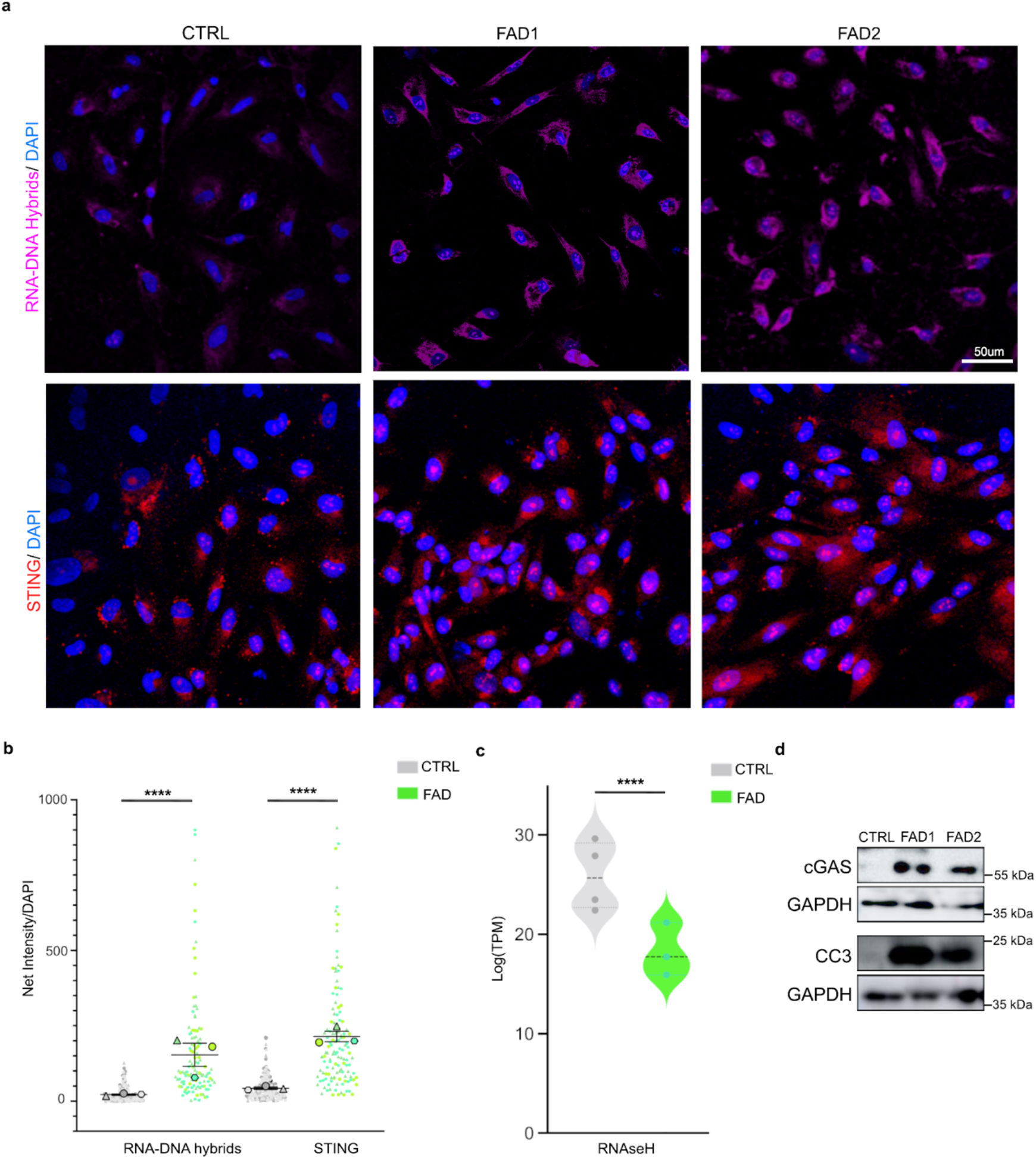
Accumulation of RNA-DNA hybrids triggers the cGAS-STING cascade and CC3 activation in FAD hpNPCs. **a**. Immunofluorescence for RNA-DNA hybrids (S9.6 antibody, pink signal) and STING (red signal) displays the accumulation of RNA-DNA hybrids in the cytoplasm of FAD hpNPCs, and an upregulation of STING. Scale bar 50um, 40X magnification. DAPI staining on nuclei in blue. **c**. Violin plot of log^2^(TPM) for RNaseH in CTRL and FAD hpNPCs. **d**. Immunoblots for cGAS, STING, and cleaved caspase 3 (CC3) in FAD hpNPCs relative to CTRLs.

These experiments provide a solid mechanistic link between aberrant TE mobilization, cytoplasmic accumulation of RNA-DNA hybrids, and cGAS-STING activation potentially resulting in cell death characterized by caspase-3 activation in AD progenitors.

### c-JUN inhibition rescues the neurogenic defects observed in the FAD hippocampal progenitors

Our genomic data revealed a distinct role for c-JUN. In fact, many of the differentially expressed genes are known c-JUN-regulated genes, and many of the DA regions harbor the JUN binding motif. Our data also shows that *JUN* is upregulated in the FAD progenitors and the activation of the WNT/JNK pathway may trigger phosphorylation of the upregulated c-JUN, leading to increased aberrant c-JUN activation. We hypothesize that these processes may lead to aberrant AP-1 activity, resulting in the opening of thousands of genomic regions harboring the JUN binding motif, allowing for the de-repression of hundreds of TEs that are typically repressed in neural precursors.

To functionally validate our genomic data and test this hypothesis, we treated CTRL and FAD hpNPCs with a synthetic peptide competitor for the binding of JNKs, called c-JUN peptide (see methods). Treatment with this peptide for five days disrupts the interaction between JNK and c-JUN, ultimately inhibiting c-JUN phosphorylation and therefore pathway activation (**Fig. 5a**).Notably, treatment of the progenitors with the c-JUN inhibitor rescues the neurogenic defects previously observed (**Fig. 1**) in the FAD progenitors. Namely, in the c-JUN inhibitor-treated FAD progenitors (hereafter FAD+c-JUN peptide), the expression of TBR2 and FOXG1 becomes comparable to the CTRLs (**Fig. 5b**), suggesting a rescue of the intermediate progenitor pool.

**Figure 5.**
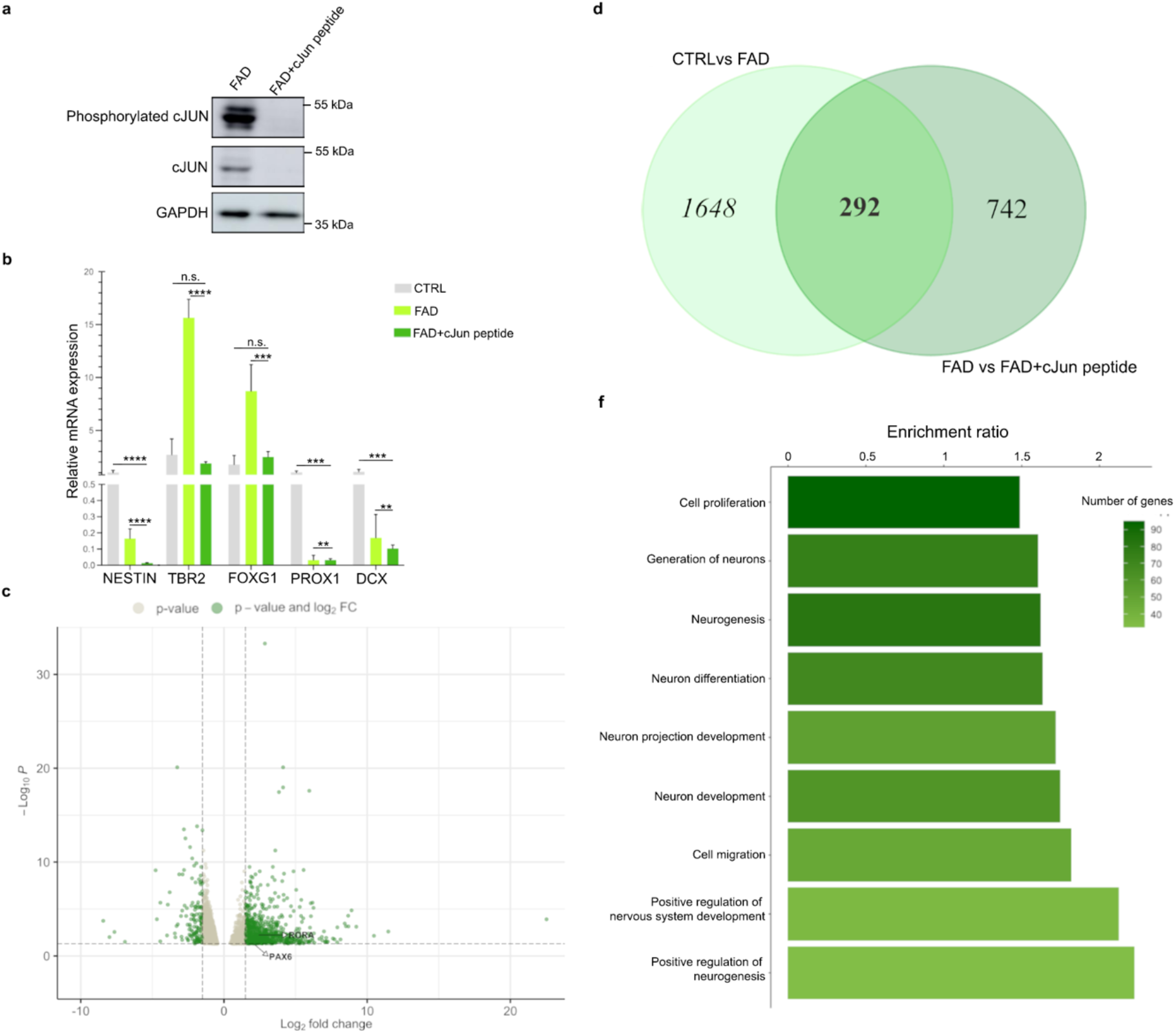
Inhibiting c-JUN phosphorylation partially rescues the impaired neurogenesis and the gene expression differences in FAD hpNPCs. **a**. Immunoblot of c-JUN and phosphorylated c-JUN untreated and treated with c-JUN peptide (FAD+c-JUN peptide) in FAD hpNPCs. **b**. qPCR analysis for hpNPCs markers in CTRL hpNPCs, untreated FAD hpNPCs, and c-JUN peptide-treated FAD hpNPCs **c**. Volcano plot of differentially expressed genes in FAD+c-JUN peptide relative to untreated FAD. Labeled genes are involved in neurogenesis (*PAX6*) and WNT/JNK signaling (*RORA*). Green = differentially expressed genes passing significance thresholds p-value < 0.05 and log^2^(fold-change) +/– 1.5; Gray = differentially expressed genes passing significance threshold of p-value < 0.05. **d**. Venn diagram displaying the genes that were differentially expressed both in the “FAD+c-JUN peptide *vs* untreated FAD” comparison and in the “FAD *vs* CTRL” comparison. 292 differentially expressed genes overlap in the two comparisons. (venn diagram was made with https://bioinformatics.psb.ugent.be) **e**. Pathways enriched in the 292 overlapping genes predicted by WebGestalt.

Next, we performed RNA-seq on FAD+cJUN peptide and untreated FAD progenitors, using both FAD lines. This experiment led to the identification of 1,034 differentially expressed genes (FDR <5%, log2(FC) = ± 1.5; **Fig. 5c**). Notably, nearly a third of these genes (292/1034) were previously identified as differentially expressed when comparing CTRL and FAD progenitors (**Fig. 5d**), suggesting that inhibiting c-JUN phosphorylation rescues a significant fraction of the transcriptomic aberrations observed in the FAD hippocampal neural precursors.

WebGestalt analysis on these 292 genes revealed enrichment for neuronal differentiation processes and inflammation (**Fig. 5e**). For 83 of the 292 genes, including the early progenitor marker *PAX6*, the expression is entirely rescued by c-JUN inhibition (**Extended data table 2**). The 83 genes with complete rescue are enriched for pathways associated with neuron differentiation (p-value = 0.00021), neuron development (p-value = 0.000324), and neuron generation (p-value = 0.000625).

These results demonstrate that c-JUN plays a crucial role in regulating neurogenesis and neuron differentiation in FAD hippocampal progenitors, and that aberrant c-JUN activity underlies a significant fraction of the transcriptomic abnormalities observed in FAD progenitors.

### c-JUN inhibition rescues TE de-repression, RNA-DNA hybrid formation, cGAS-STING activation, and caspase-3 activation

c-JUN inhibition also impacts the observed aberrant opening of chromatin at TE loci. By performing an RT-qPCR on a group of RTEs, selected among those previously identified as aberrantly active in FAD, we demonstrated that c-JUN inhibition results in a significant reduction of TE transcription at both LTR and LINE loci (**Fig. 6a**).

**Figure 6.**
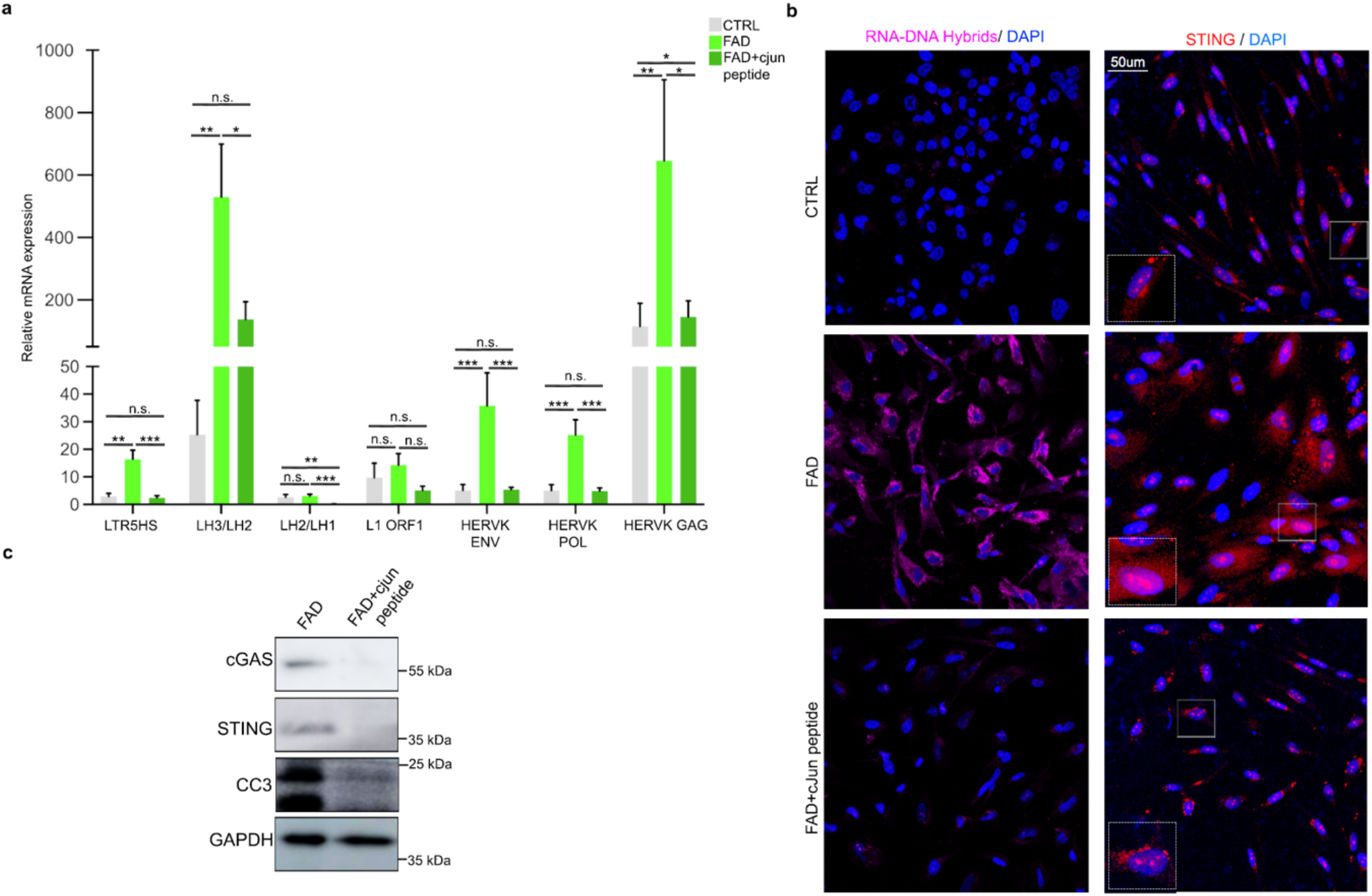
Inhibiting c-JUN phosphorylation rescues aberrant TE derepression and the activation of the cGAS-STING cascade in FAD progenitors. **a**. qPCR analysis for a group of TEs selected among those previously identified as aberrantly active in FAD hpNPC. Primers for HERVK/LTR5HS target individual ORFs from the LTR; Primers for L1-ORF1 target a conserved region in ORF1 of 6x; the other L1 primers target the L1PA2 family; the LH2/LH3 primers target the end of the 3’ UTR; The LH1/LH2 primers target the 5’ of the other amplicon. **b**. Immunofluorescence for RNA-DNA hybrids (S9.6 antibody, pink signal) and STING (red signal) shows that c-JUN inhibition significantly decreases the accumulation of RNA-DNA hybrids and STING levels in the cytoplasm of FAD hpNPCs. Scale bar 50um, 40X magnification. White dot line boxes represent 2X magnification of the corresponding squared box. DAPI staining on nuclei in blue. **c**. Immunoblots for cGAS, STING, and cleaved caspase 3 (CC3) on FAD+c-JUN peptide hpNPCs relative to FAD progenitors.

Next, we tested if the reduction of aberrant TE mobilization, resulting from c-JUN inhibition (**Extended Data Fig. 3a**) affected RNA-DNA dimer accumulation and consequent cGAS-STING activation. Notably, FAD+cJUN peptide progenitors show a significant reduction of both RNA-DNA hybrid accumulation and cGAS-STING cascade activation relative to untreated cells (**Fig. 6b, c**; **Extended Data Fig. 3b**). Finally, an immunoblot for cleaved caspase 3 indicated that inhibition of the JNK/c-JUN interaction also prevented activation of CC3 in FAD progenitors (**Fig. 6c**).

With these results, we demonstrate that by inhibiting c-JUN phosphorylation, the chromatin remains in a repressed state. Therefore, aberrant TE mobilization does not occur, preventing the activation of the cGAS–STING-induced caspase-3 activation.

### Aberrant c-JUN activity underlies cGAS-STING activation and caspase-3 activation in sporadic AD hippocampal progenitors

To test whether the mechanism that we characterized in FAD progenitors is shared between the two AD types (familial and sporadic) we differentiated two sporadic Alzheimer’s iPSC lines (SAD1 and SAD2) into hpNPCs, following the same protocol previously described (**Extended Data Fig. 4a**). Notably, an RT-qPCR for the same markers characterized for FAD (NESTIN, TBR2, FOXG1, PROX1, and DCX) also revealed impaired neurogenesis in SAD, with an accelerated differentiation signature consistent with previous studies in sporadic AD (**Extended Data Fig 4b**)^77^. Interestingly, RNA-seq revealed that *JUN* is also upregulated in the SAD progenitors (**Extended Data Fig 4c**) and that the 183 differentially expressed genes in SAD hpNPCs are involved in neurogenesis and neuronal differentiation pathways (Extended Data Fig 5d). Moreover, as in FAD progenitors, SAD hpNPCs show abnormal chromatin accessibility at TE loci (**Extended Data Fig 4e**).

Treatment of SAD progenitors with the inhibitor of c-JUN phosphorylation led to a decrease in cytoplasmic RNA-DNA dimers (**Fig. 7a**), as well as a significant a reduction of cGAS/STING and cleaved caspase 3 activation (**Fig. 7b**). Importantly, these experiments revealed that both AD types (familial and sporadic) share the same aberrantly activated cell-death axis.

**Figure 7.**
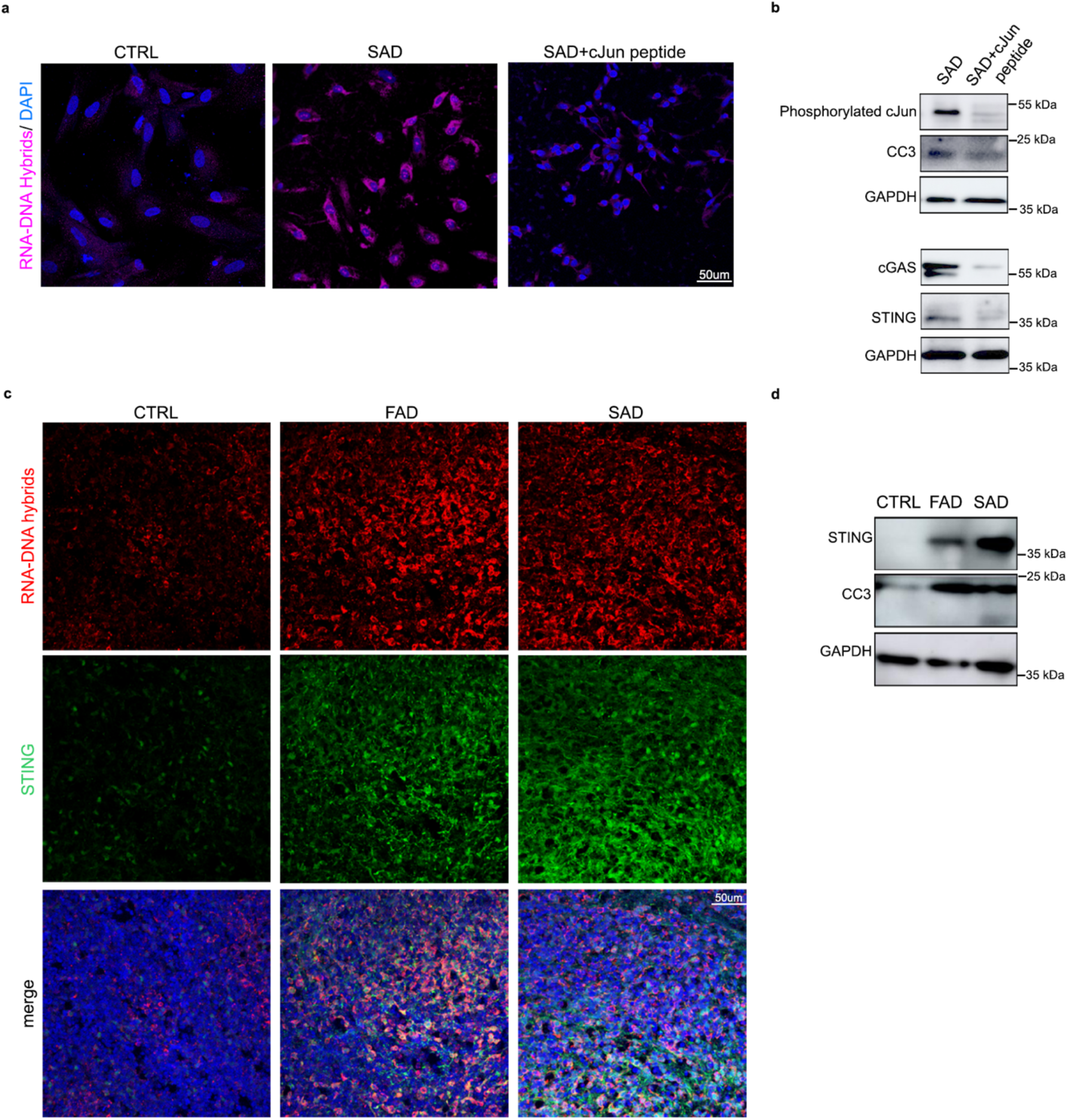
TE-derived RNA-DNA hybrids and cGAS-STING activated in SAD progenitors and AD cerebral organoids. **a**. Immunofluorescence for RNA-DNA hybrids (S9.6 antibody, pink signal) and STING (red signal) show that c-JUN inhibition significantly decreases the accumulation of RNA-DNA hybrids and STING levels in the cytoplasm of SAD hpNPCs. Scale bar 50um, 40X magnification. DAPI staining on nuclei in blue. **b**. Immunoblots for phosphorylated c-JUN, cGAS, STING, and cleaved caspase 3 (CC3) on SAD+c-JUN peptide hpNPCs relative to SAD progenitors. **c**. Immunofluorescence for RNA-DNA hybrids (S9.6 antibody, red signal) and STING (green signal) on cerebral organoids (CTRL, FAD, SAD). Scale bar 50um, 40X magnification. DAPI staining on nuclei in blue. **d**. Immunoblots for STING and cleaved caspase 3 (CC3) on AD organoids relative to CTRLs.

### RTE-derived RNA-DNA hybrids inducing the cGAS-STING cell-death axis are present in familial and sporadic AD cerebral organoids

Finally, we tested whether this pathway was also active in cerebral organoids, which harbor both progenitor cells and differentiated neurons in a three-dimentional architecture, resembling the human brain. The FAD, SAD, and CTRL iPSC lines were differentiated into cerebral organoids through an embryoid body intermediate. After 62 days, immunofluorescence was performed on cerebral organoids exhibiting the proper neuronal differentiation (**Extended Data Fig. 5a**). We observed a more significant overall accumulation of RNA-DNA hybrids and an upregulation of STING in FAD and SAD organoids compared to CTRL organoids (**Fig. 7c, Extended Data Fig. 5b**). The activation of the cGAS-STING cell-death axis and the increase in CC3 were also confirmed through western blot (**Fig. 7d**). As expected, the cytoplasmic accumulation of RNA-DNA hybrids was seen in TBR2-positive neural progenitors (**Extended Data Fig 5c**) that are enriched in the FAD organoids (**Extended Data Fig 5c**). Interestingly, mature neurons (MAP2-positive) in AD cerebral orgnaoids also have a cytoplasmatic accumulation of RNA-DNA hybrids and an upregulation of STING (**Extended Data Fig 5d**). Finally, activating the cGAS-STING cell-death axis in neurons leads to caspase-3 activation (**Extended Data Fig 5e**). These organoid-based data further validated our proposed pathogenic mechanism and cascade for both familial and sporadic AD. This suggests that the molecular impairment driven by TE re-activation observed in progenitors, is also maintained in mature neurons in a physiological model of neural differentiation.

Overall, this collection of experiments unveils a novel mechanism linking AP-1 to TE mobilization, innate immunity and cell death in AD.

## Discussion

Alzheimer’s disease (AD) is the most common neurodegenerative disorder. Fibrillar deposits of highly phosphorylated TAU protein are a key pathological feature of AD as well as other AD related dementias^78^. Importantly, studies in *Drosophila* have highlighted that TAU hyperphosphorylation and upregulation correlate with global nuclear chromatin relaxation and abnormal transcriptional activation of heterochromatic genomic regions^40,43^. One of the consequences of this phenomenon is the aberrant mobilization of TEs, which are typically repressed in the genome. Progressive TE de-repression and mobilization in the brain typically correlates with ageing^39,44^. However this phenomenon is significantly exacerbated in many neurodegenerative disorders, including Amyotrophic Lateral Sclerosis (ALS), Multiple Sclerosis, and Alzheimer’s^39,41–46^.

Recent studies showed that overexpression of TAU alone, in aging *Drosophila* brains is sufficient to increase the expression of retrotransposons, mostly belonging to the LINE and ERV groups^40,43^. However, the mechanism linking tauopathies to chromatin relaxation and TE mobilization remains unexplored.

Here, we unveiled a cascade of biological processes which links the upregulation of the AP-1 member, c-JUN, to aberrant TE mobilization. We detected an upregulation of c-JUN in hippocampal progenitors derived from both familial and sporadic AD iPSC lines. Further, kinases involved in the regulation and phosphorylation of c-JUN (MAPK/JNK signaling) were also dysregulated in fully differentiated FAD iPSC-derived CA3 hippocampal neurons. We demonstrated that c-JUN upregulation has two main consequences: 1) the dysregulation of hundreds of genes involved in neuronal differentiation and neuron generation 2) the activation of hundreds of RTEs, that harbor the AP-1 binding motif. AP-1 activating repressed chromatin was not an unexpected result; several recent studies have demonstrated that AP-1 can act as a pioneer transcription factor by recruiting the BAF chromatin remodeling complex to its targets to elicit accessibility and activation^35,36,79^.

The second goal of our study was to investigate the consequences of aberrant TE de-repression and mobilization. With our experiments, we demonstrated that abnormal expression of RTEs leads to the accumulation of RTE-derived RNA-DNA hybrids in the cytoplasm of the AD hpNPCs, as well as in AD cerebral organoids. We showed that this triggers the activation of the innate immune response, particularly of the cGAS-STING pathway, which ultimately elicits the accumulation of cleaved caspase-3, a molecular signature of cells undergoing apoptosis^80^. Importantly, we observed this phenomenon in both familial and sporadic AD lines. We were able to explicitly demonstrate that c-JUN facilitates this mechanism and that treating the AD hpNPCs with a c-JUN inhibitor sufficiently decreases this cascade leading to a reduction of cell death.

Neuroinflammation plays an important role in the pathogenesis of Alzheimer’s disease, and in this context, the cGAS–STING signalling pathway has recently emerged as a key mediator of inflammation in the settings of infection, cellular stress and tissue damage.^81^ In fact, neuroinflammation is primarily driven by type-I interferons (INFs), and the role of STING in the control of the type-I IFN-mediated mediated response is becoming increasingly appreciated.^82^ Given this premise, the finding of our studies may open new potential therapeutic avenues. For instance, nanobody-based targeting^83^ of cytoplasmatic RNA-DNA hybrids might provide a new therapeutic approach to counteract the activation of innate immune response and to reduce neuroinflammation, bypassing the side effects of the anti-inflammatory drugs currently involved in the clinical trials. Additionally, experiments on pre-clinical models may be employed to test compounds which could act downstream in the cell death axis demonstrated here. Finally, the cytoplasmic accumulation of RNA-DNA hybrids could be used as an early biomarker for AD in imaging tools for diagnosis.

In summary, these lines of evidence point toward a pathological mechanism underlying AD. Future studies on possible therapeutic intervention that would target this mechanism are essential with the goal of identifying therapeutic strategies and early diagnosis for AD and other neurogenerative disorders.

## Acknowledgements

The authors are thankful to Julius Judd and Cedric Feschotte (Cornell) for sharing the primers for RT-PCR at TE loci, and for insightful discussion of the findings. The authors thank the Jefferson Stem Cell Center for preliminary expansion and characterization of the patient lines, and for helping with the optimization of the differentiation protocols. This study was funded by NIH R35 GM138344-01, awarded to M.T. The following sources also supported this work: National Institutes of Health R21-NS090912 (D.T.), RF1-AG057882 (D.T.), Muscular Dystrophy Association grant 628389 (D.T.); DoD grant W81XWH-21-1-0134 (M.E.C.); and Farber Family Foundation (D.T., C.S.).

## Data availability

The original genome-wide data generated in this study have been deposited in the GEO database under accession code GSE213610.

## Author contributions

M.T., D.T and C.S. designed the experiments. C.S. performed most of the experiments. S.M.B, performed some of the experiments. M.E.C and M.S. contributed to some experiments and analyses. C.S., M.T., S.M.B. and D.T. analyzed the data. C.S. wrote the manuscript with the contribution of all the authors.

## Materials and Methods

### Human iPSC culture

Control and Alzheimer’s disease iPSC lines were obtained from the Coriell Institute for Medical Research (Camden, NJ). In particular, we received two control lines (Control line-1:IPSM8Sev3, male, 65 years old and Control line-2: iPSM15Sev4, female, 62 years old) CTRL1 and CTRL2 respectively; two familial Alzheimer’s disease lines (Familial Alzheimer’s line-1: AG25370, female, 80 years old and Familiar Alzheimer’s line-2: GM24675, male, 60 years old) FAD1 and FAD2 respectively; and two sporadic Alzheimer’s disease lines (Sporadic Alzheimer’s line-1: AG27607, female, 69 years old and Sporadic Alzheimer’s line-2: GM24666, male, 83 years old) SAD1 and SAD2 respectively. All the AD lines excepted for AG25370 were validated in previous studies^77,83,85,86^.

The iPSC lines were expanded in feeder-free, serum-free mTeSR™1 medium (85850, STEMCELL Technologies). Cells were passaged ∼1:10 at 80% confluency using EDTA 0.5mM (15575020, Invitrogen) and small cell clusters (50–200 cells) were subsequently plated on tissue culture dishes coated overnight with Geltrex™ LDEV-Free hESC-qualified Reduced Growth Factor Basement Membrane Matrix (A1413302, Fisher-Scientific).

### hpNPC Differentiation

The iPSC lines were differentiated into hpNPCs as previously described^47^. Briefly, iPSCs were treated with hpNPC induction medium for five days: DMEM/F-12 medium (Invitrogen) supplemented with B-27 (A3582801, Gibco), N-2 (17502048, Gibco), DKK1 (778606, Biolegend), Cyclopamine (C-8700, LC Laboratories), Noggin (597004, Biolegend), and SB431542 (S1067, Selleck Chemicals LLC). At day 6, the hpNPCs were plated in a new geltrex-coated well and cultured in proliferation medium, consisting of DMEM/F-12 medium (Invitrogen) supplemented with B-27 (A3582801, Gibco), N-2 (17502048, Gibco) and 20 ng/ml bFGF (713304, Biolegend).

### CA3 Neuron Differentiation

The iPSC lines were differentiated into CA3 Neurons as previously described^68^. Briefly, iPSCs were treated with hpNPC induction medium for 15 days. At day 16, the hpNPCs were plated in a new PLO-Laminin double-coated well in Neuron induction medium, consisting in DMEM/F12 medium (11320082, Gibco) supplemented with B-27 (A3582801, Gibco), N-2 (17502048, Gibco), BDNF (450-02, Prepotech), Dibutyryl-cAMP (11-415-0, Tocris), laminin (23017015, Thermofisher Scientific), AA (A4544-25G, Sigma), WNT3a (5036-WN, R&D System). After 3 weeks, the neurons were switched to neuron medium, consisting of DMEM/F-12 medium (Invitrogen) supplemented with B-27 (A3582801, Gibco), N-2 (17502048, Gibco), BDNF (450-02, Prepotech), Dibutyryl-cAMP (11-415-0, Tocris), laminin (23017015, Thermofisher Scientific) and AA (A4544-25G, Sigma) for one week. Mature CA3 neurons were then collected for RNA-seq and fixed for immunofluorescence.

### Cerebral organoid Differentiation

CTRL and AD (FAD and SAD) iPSCs were differentiated into cerebral organoids following a previously published protocol^86^. Briefly, embryoid bodies were formed from CTRL, FAD, and SAD iPSCs and maintained in Essential 8 media (E8 media, A1517001, Thermoscientific) supplemented with ROCK inhibitor (SCM075, Millipore) for 4 days. Neuronal induction was obtained by replacing the E8 media with Neural induction media: DMEM/F12 (11330-032, Invitrogen) supplemented with N-2 (17502048, Gibco), 1% Glutamax (35050-038, Invitrogen), 1% MEM-NEAA (M7145, Sigma), and Heparin at a final concentration of 1 μg/ml (H3149, Sigma). After 4-5 days, when neuroepithelium formation was achieved, spheroids were embedded in Matrigel (356234, BD Biosciences) and cultured in Cerebral organoid differentiation media: DMEM/F12 (11330-032, Invitrogen) and Neurobasal (21103049, Invitrogen) (1:1) supplemented with B27 without VitA (12587010, Invitrogen), N2 (17502048, Gibco), Insulin (I9278-5ML, Sigma), 2-Mercaptoethanol (1:100 dilution, 8057400005, Merk), 1% MEM-NEAA (M7145, Sigma), and Glutamax (35050-038, Invitrogen). After 4 days in static culture, spheroids were transferred to a shaker and maintained. Half media changes were performed every 3-4 days.

### hpNPC c-JUN peptide treatment

CTRL, FAD, and SAD hpNPCs were treated with 100µM of c-JUN peptide (19-891, Fisher Scientific) for 5 days in a proliferative condition. This peptide comprises residues 33–57 of the JNK binding (δ) domain of human c-JUN and it is a competitive inhibitor of JNK/c-JUN interaction preventing c-JUN phosphorylation and activation.

### Processing of organoids

At day 62, the whole organoids were fixed in 4% PFA overnight at 4 °C. After cryoprotection in 30% sucrose (s7903, Sigma), organoids were cryo-sectioned at 20 μm thickness and slices were analyzed by immunohistochemistry.

### Immunofluorescence

Immunohistochemistry of iPSCs, hpNPCs, and CA3 neurons was performed in µ-Slide 4 Well Glass Bottom (80426, IBIDI), while organoid IF was performed on 20-μm serial sections. Upon fixation (4% PFA for 10 minutes), cells were permeabilized in blocking solution (0.1% Triton X-100, 1X PBS, 5% normal donkey serum) and then incubated with the antibody of interest. The total number of cells in each field was determined by counterstaining cell nuclei with 4,6-diamidine-2-phenylindole dihydrochloride (DAPI; Sigma-Aldrich; 50 mg/ml in PBS for 15 min at RT). To improve the efficiency of Tbr2 detection, the cells and the organoid slides, prior to permeabilization and blocking step, were treated with 10 mM sodium citrate (pH = 6) for 10 minutes at 95 °C.

For RNA-DNA hybrid staining (S9.6 antibody), upon fixation (4% PFA for 10 minutes), cells and organoid slides were permeabilized in PBS 1X 0.5% Triton X-100 for 15 minutes. They were then incubated overnight at -20°C in 100% methanol. The samples were then blocked in 1X PBS 5% NDS for 4 hours at 37°C and followed by overnight incubation with the S9.6 antibody.

Immunostained cells and organoid slices were analyzed via confocal microscopy using a Nikon A1R+. Images were captured with x40 for hpNPCs and x20 and x60 objectives for organoids and a pinhole of 1.0 Airy unit. Analyses were performed in sequential scanning mode to rule out cross-bleeding between channels. Fluorescence intensity quantification was performed with Fiji and the NIS-Elements AR software. All antibodies are listed in the Antibodies table (Supplementary Table S3).

### Western Blot

For total lysate, cells were harvested and washed three times in 1X PBS and lysed in RIPA buffer (50mM Tris-HCl pH7.5, 150mM NaCl, 1% Igepal, 0.5% sodium deoxycholate, 0.1% SDS, 500uM DTT) with protease and phosphatase inhibitors. Twenty μg of whole cell lysate were loaded in Novex WedgeWell 4-12% Tris-Glycine Gel (Invitrogen) and separated through gel electrophoresis (SDS-PAGE) in Tris-Glycine-SDS running buffer (Invitrogen). The proteins were then transferred to ImmunBlot PVDF membranes (ThermoFisher) for antibody probing. Membranes were incubated with 10% BSA in 1X TBST for 1 hour at room temperature (RT), then incubated for variable times and concentrations with the suitable antibodies (Supplementary table S3) diluted in 5% BSA in 1X TBST. Membranes were then washed with 1X TBST and incubated in the HRP-linked species-specific secondary antibody (1:10000 dilution) for one hour at RT. The membrane was visualized using the Pierce ECL Plus Western Blotting Substrate (32132; ThermoFisher) and imaged with an Amersham Imager 680. All antibodies are listed in the Antibodies table (Supplementary Table S3).

### Real-time quantitative polymerase chain reaction (RT-qPCR)

Cells were lysed in Tri-reagent (R2050-1-50, Zymo Research) and RNA was extracted using the Direct-zol RNA Miniprep kit (Zymo Research). 600ng of template RNA was retrotranscribed into cDNA using RevertAid first strand cDNA synthesis kit (Thermo Scientific) according to the manufacturer’s directions. 15ng of cDNA was used for each real-time quantitative PCR reaction with 0.1 μM of each forward and reverse primer, 10 μL of PowerUp™ SYBR™ Green Master Mix (Applied Biosystems) in a final volume of 20 μl, using a QuantStudio 3 Real-Time PCR System (Applied Biosystems). Thermal cycling parameters were set as follows: 3 minutes at 95°C, followed by 40 cycles of 10 seconds at 95°C and 20 seconds at 63°C followed by 30 seconds at 72°C. Each sample was run in triplicate. 18S rRNA was used for normalization. Primer sequences are reported in Supplementary Table S4.

### RNA-Seq

Cells were lysed in Tri-reagent (R2050-1-50, Zymo Research) and total RNA was extracted using Quick-RNA Miniprep kit (R1055, Zymo Research) according to the manufacturer’s instructions. RNA was quantified using a DeNovix DS-11 Spectrophotometer while the RNA integrity number (RIN) was checked on an Agilent 2200 TapeStation. Only samples with RIN values above 8.0 were used for transcriptome analysis. RNA libraries were prepared using NEBNext^®^ Poly(A) mRNA Magnetic Isolation Module (E7490S, New England Biolabs), NEBNext^®^ UltraTM II Directional RNA Library Prep Kit for Illumina^®^ (E7760S, New England Biolabs) and NEBNext^®^ UltraTM II DNA Library Prep Kit for Illumina^®^ (E7645S, New England Biolabs) according to the manufacturer’s instructions. The libraries were sequenced using an Illumina NextSeq2000, generating 150 bp Paired-End reads.

### RNA-Seq Analyses

Reads were aligned to hg19 using STAR v2.5^87^ in 2-pass mode with the following parameters: --quantMode TranscriptomeSAM --outFilterMultimapNmax 10 --outFilterMismatchNmax 10 --outFilterMismatchNoverLmax 0.3 --alignIntronMin 21 -- alignIntronMax 0 -- alignMatesGapMax 0 --alignSJoverhangMin 5 --runThreadN 12 -- twopassMode Basic -- twopass1readsN 60000000 --sjdbOverhang 100. We filtered bam files based on alignment quality (q = 10) using Samtools v0.1.19 (Li H, et al., 2009). We used the latest annotations obtained from Ensembl to build reference indexes for the STAR alignment. Kallisto (Bray et al., 2016) was used to count reads mapping to each gene. RSEM^88^ was used to obtain FPKM (Fragments Per Kilobase of exon per Million fragments mapped). Differential gene expression levels were analyzed using DESeq2^89^, with the following model: design = ∼condition, where condition indicates either CTRL or Alzheimer’s disease (FAD or SAD) lines.

### ATAC-Seq

For ATAC-Seq experiments, 50,000 cells per condition were processed as described in the original ATAC-seq protocol paper^90^. Briefly, 50,000 cells were collected, washed, and lysed. The chromatin was subjected to transposition/library preparation via a Tn5 transposase using the Tagment DNA Enzyme and Buffer Kit (20034197, Ilumina) and incubated at 37°C for 30 min with slight rotation (300 RPM). Transposed DNA was purified using a MinElute PCR Purification Kit (28004; Qiagen). Transposed DNA fragments were then amplified using a universal and barcoded primer^90^. Thermal cycling parameters were set as follows: 1 cycle of 72°C for 5 minutes, 98°C for 30 seconds, followed by 5 cycles of 98°C for 10 seconds, 63°C for 30 seconds, and 72°C for 1 min. The amplification was paused and 5ul of the partially amplified, transposed DNA was used for a qPCR side reaction including the universal and sample-specific barcoded primers^90^, PowerUp™ SYBR™ Green Master Mix (Applied Biosystems), NEBNext High-Fidelity 2x PCR Master Mix, and nuclease-free water. The qPCR side reaction parameters were set as follows: 1 cycle of 72°C for 5 minutes, 98°C for 30 seconds, followed by 40 cycles of 98°C for 10 seconds, 63°C for 30 seconds, and 72°C for 1 min. The Rn vs cycle plot was used to determine the remaining number of PCR cycles needed where 1/3 of the maximum fluorescent intensity corresponds to the cycle number. The remaining partially amplified transposed DNA was fully amplified using the previous parameters with the additional cycle number determined from the qPCR side reaction. The amplified, transposed DNA was purified using AMPure XP beads (A63881, Beckman Coulter) and sequenced using an Illumina NextSeq2000, generating 150 bp Paired-End reads.

### ATAC-Seq Analyses

After removing the adapters, the sequences were aligned to the reference hg19, using Burrows-Wheeler Alignment tool (BWA), with the MEM algorithm^91^. Aligned reads were filtered based on mapping quality (MAPQ > 10) to restrict our analysis to higher quality and likely uniquely mapped reads, and PCR duplicates were removed. All mapped reads were offset by +4 bp for the forward strand and -5 bp for the reverse strand. We called peaks using MACS2^92^, at 5% FDR, with default parameters. We analyzed differential genome accessibility using DESeq2^89^, with the following model: design = ∼condition, where condition indicates either CTRL or Alzheimer’s disease (FAD or SAD) lines. R v3.3.1. and BEDtools v2.27.1^93^ were used for all comparative TEs analyses.

### Statistical and genomic analyses

All statistical analyses were performed using R v3.3.1. BEDtools v2.27.1^93^ was used for genomic studies. Pathway analysis was performed with WEB-based GEne SeT AnaLysis Toolkit (http://www.webgestalt.org). Motif analyses were performed using the MEME-Suite^95^, specifically with the MEME-ChIP application. Fasta files of the regions of interest were produced using BEDTools v2.27.1. Shuffled input sequences were used as background. E-values < 0.001 were used the threshold for significance. All described results (qPCR analyses and Immunofluorescences) are representative of at least three independent experiments unless specifically stated otherwise. Data were presented as average ± SEM. Statistical analysis was performed using Excel (Microsoft) or GraphPad Prism 8 software (GraphPad). Student’s t-test was used for the comparison between two groups. A value of P < 0.05 was considered significant; *P < 0.05; **P < 0.01; ***P < 0.001; ****P < 0.0001; n.s., not significant.

## Extended Data

**Extended Data Figure 1.**
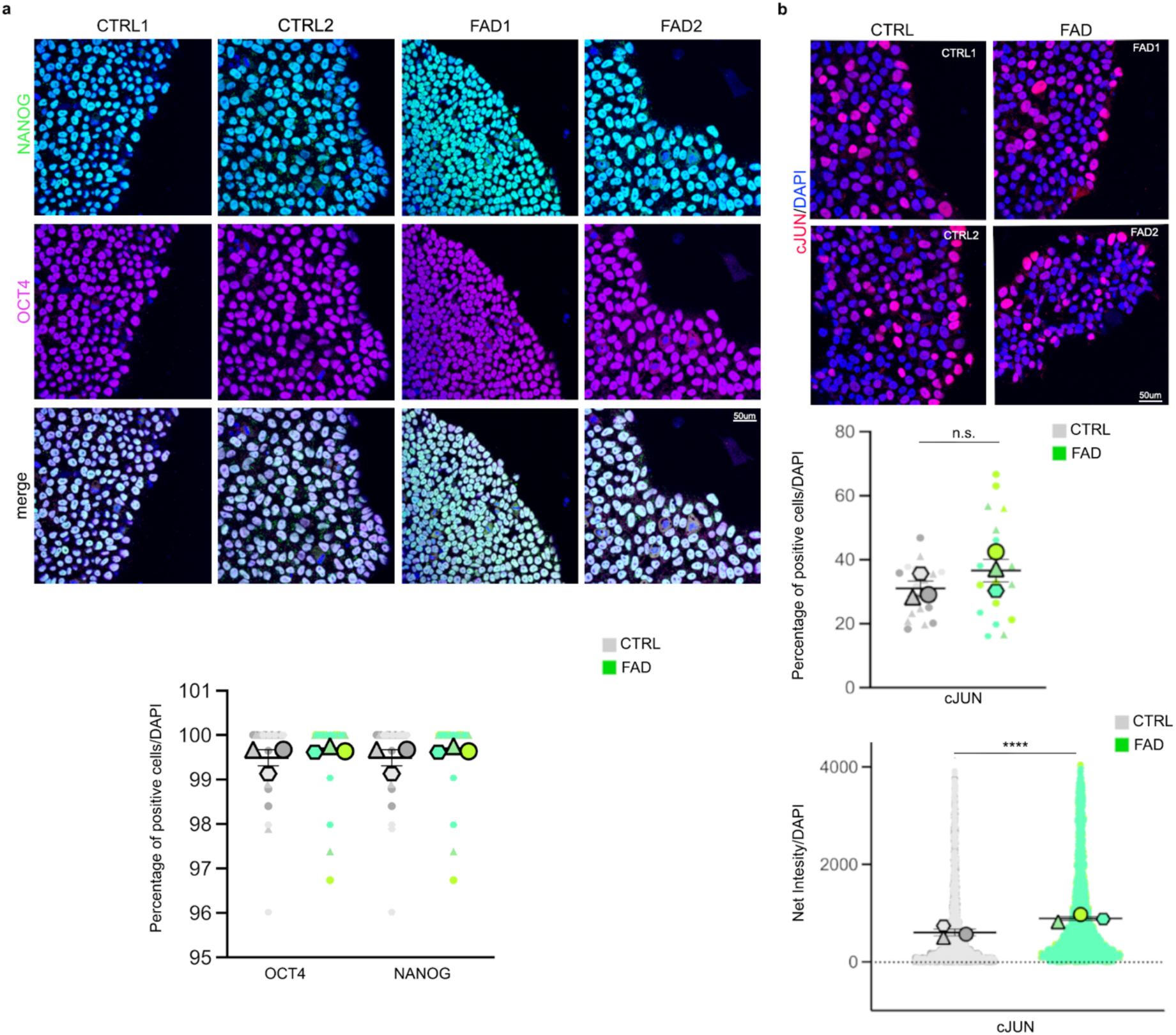
Pluripotency and JUN expression profiles of CTRL and FAD-derived iPSCs. **a**. and **b**. Immunofluorescence quantifying the expression of **a**. pluripotency markers OCT4 (purple) and NANOG (green) and **b**. endogenous c-JUN in iPSCs derived from CTRLs and FAD. The quantification of immunofluorescence in panel b is reported as the percentage of expressing cells (upper superplot) and expression of Net Intesity (bottom superplot).

**Extended Data Figure 2.**
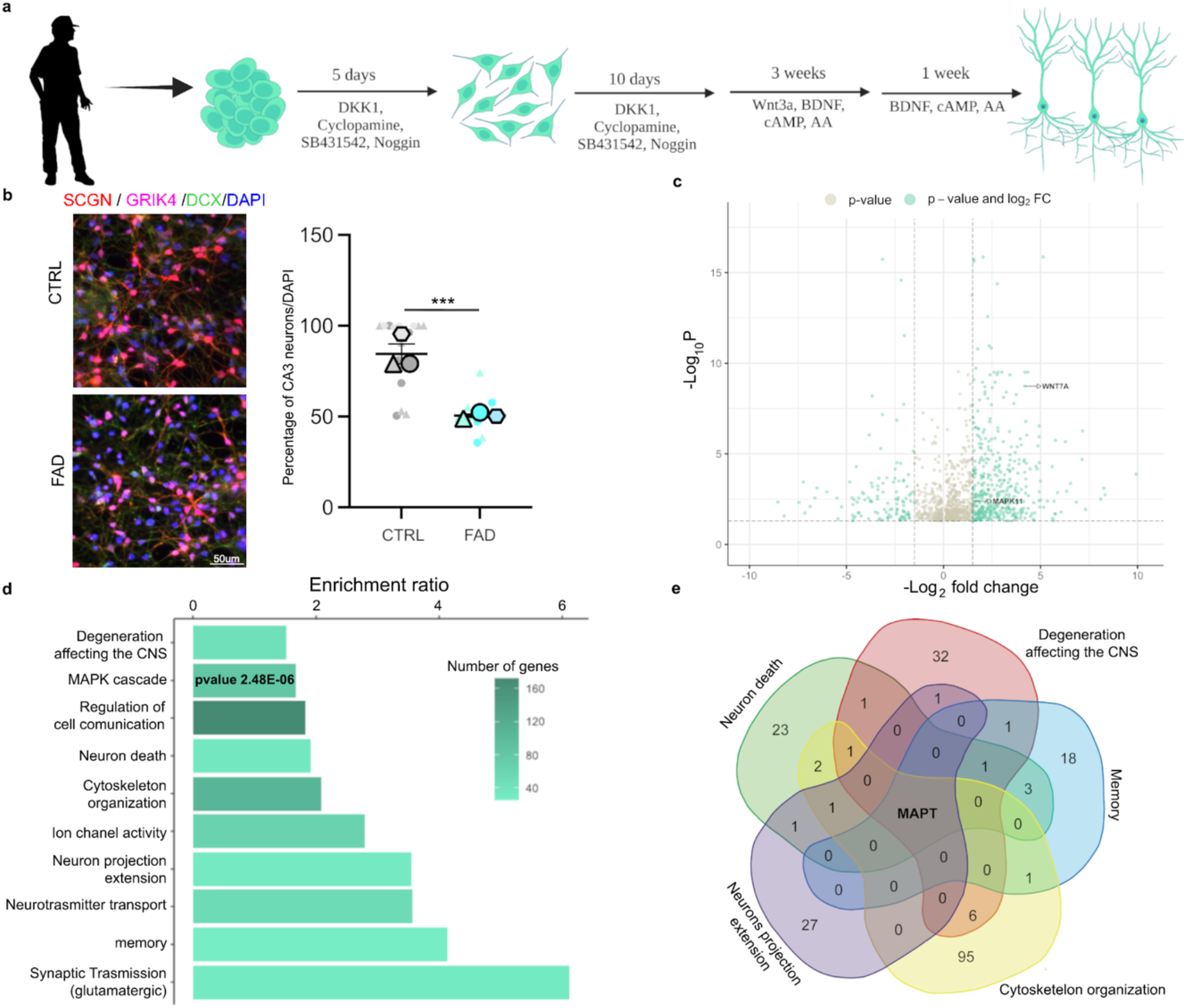
Dysregulated transcriptional networks and pathways in FAD CA3 hippocampal neurons. **a**. A scheme of the protocol for CA3 hippocampal neuron differentiation (made with Biorender.com). **b**. Immunofluorescence for CA3 neuron markers. GRIK4 (pink) and SCGN (red) were expressed in CTRL and FAD neurons demonstrating their proper differentiation. The relative quantification shows that FAD hpNPCs failed to differentiate into CA3 neurons as demonstrated by the reduced number of SCGN- and GRIK4-positive cells in the FAD neuron culture. Scale bar 50um, 40X magnification. DAPI staining on nuclei in blue. **c**. Volcano plot displaying the 1,105 differentially expressed genes in FAD CA3 neurons relative to CTRL CA3 neurons. Teal = differentially expressed genes passing significance thresholds p-value < 0.05 and log^2^(fold-change) +/– 1.5; Gray = differentially expressed genes passing significance threshold of p-value < 0.05. **d**. Enriched pathways associated with the 1,105 differentially expressed genes in FAD CA3 neurons predicted by WebGestalt. **e**. Venn diagram showing the genes shared across five enriched pathways in AD. (venn diagram was made using https://bioinformatics.psb.ugent.be).

**Extended Data Figure 3.**
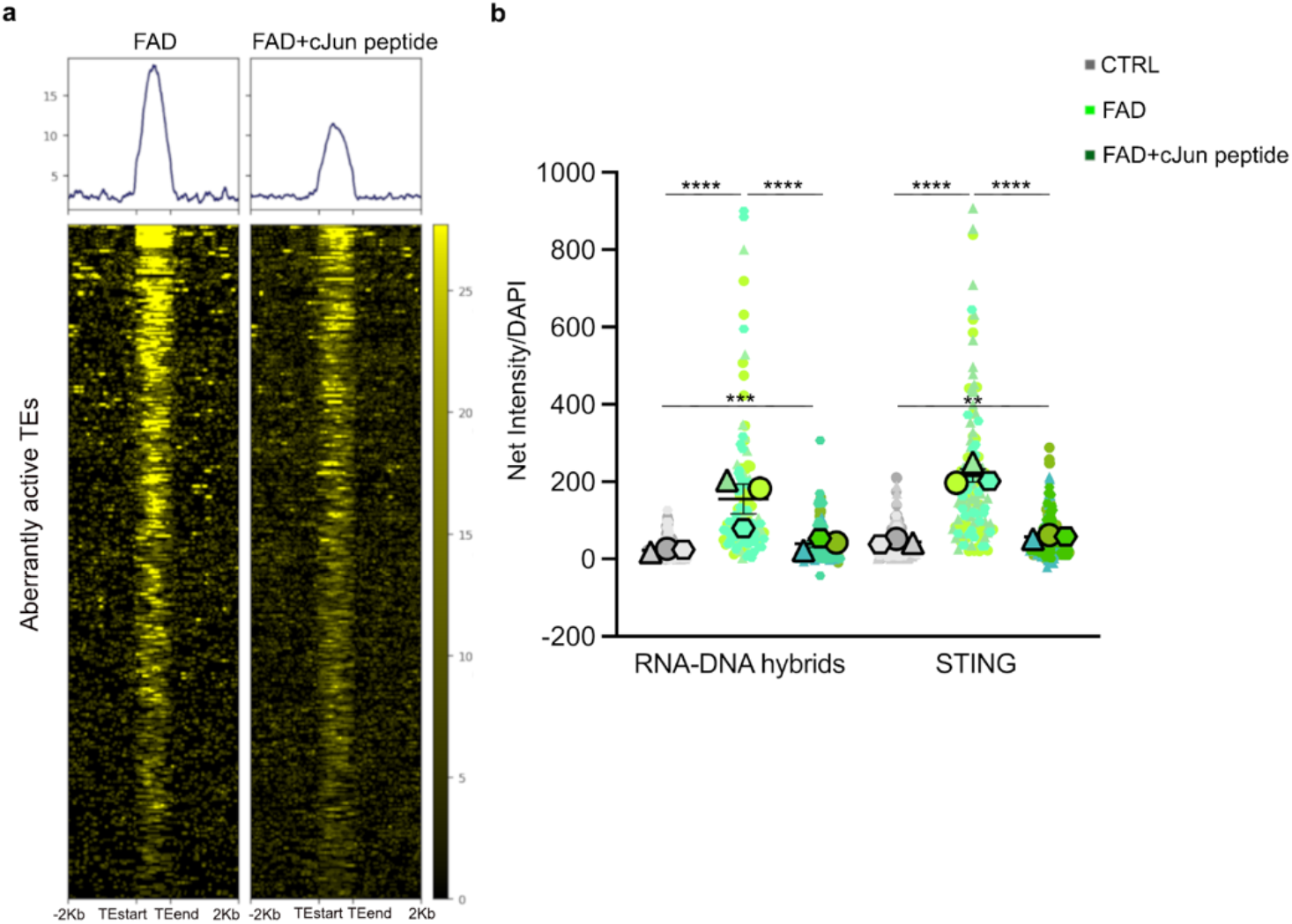
TE activation induces RNA-DNA hybrid accumulation triggering the cGAS-STING cascade and apoptosis in FAD hpNPCs. **a**. Heatmap showing a reduction in the number of open regions of chromatin in FAD+c-JUN peptide at aberrantly active TEs. **b**. Quantification of the immunofluorescences in Figure 4a.

**Extended Data Figure 4.**
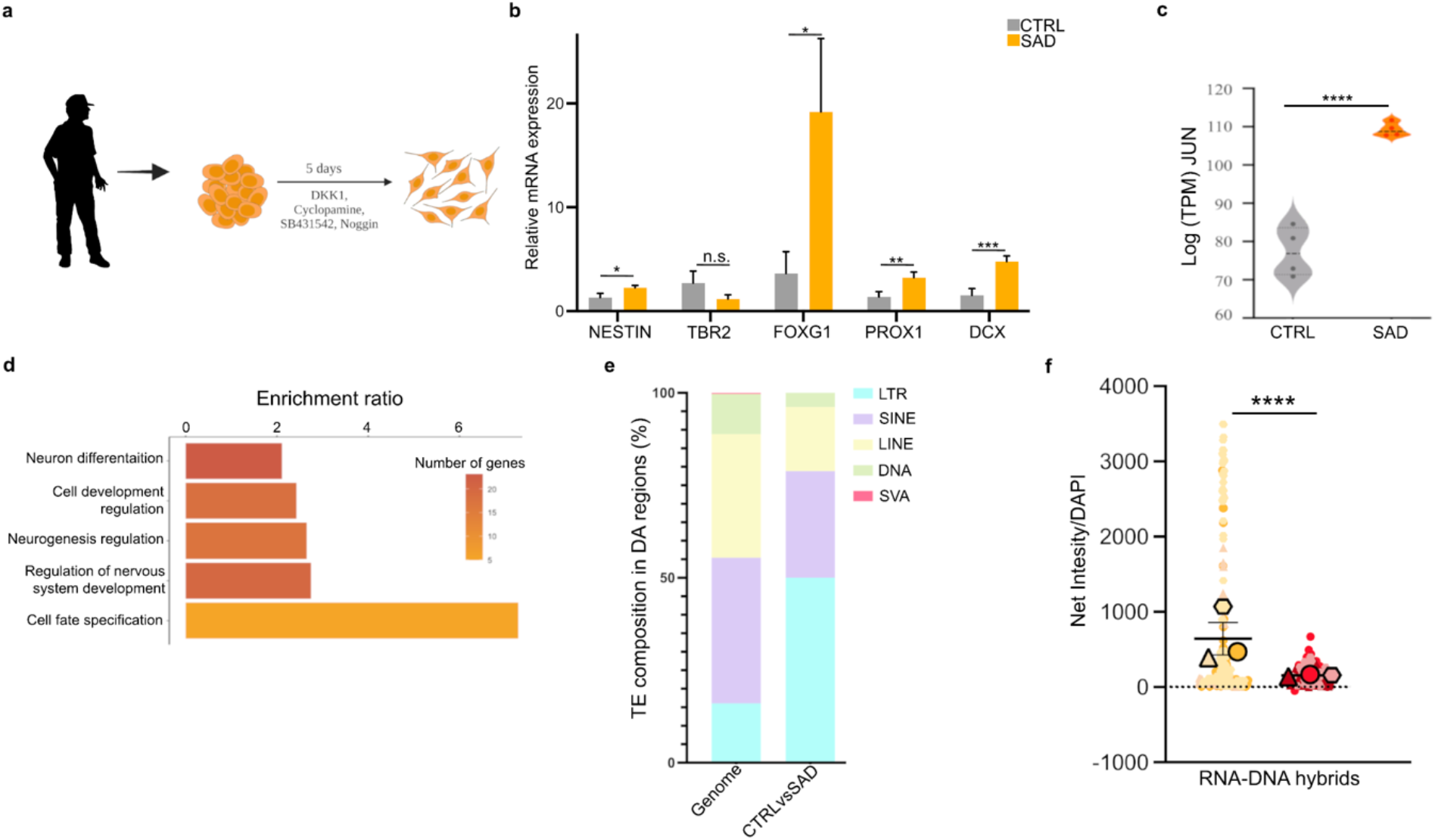
SAD iPSC-derived hippocampal neural progenitors display impaired neurogenesis, gene expression dysregulation, and aberrant activation of TEs. **a**. The scheme of the protocol for hpNPC differentiation (made with Biorender.com). **b**. qPCR analysis for hpNPC population markers. NESTIN – early precursors; TBR2/FOXG1 – intermediate progenitors; PROX1 – late progenitors; DCX – neuroblasts. **c**. Violin plot of log^2^(TMP) for *JUN* in SAD hpNPCs compared to CTRLs. **d**. Pathways enriched in the 183 differentially expressed genes (SAD vs CTRL hpNPCs) predicted by WebGestalt. **e**. TE family distribution of the TEs aberrantly de-repressed in SAD progenitors shows an enrichment for LTRs. **f**. Quantification of the immunofluorescences in Figure 7a.

**Extended Data Figure 5.**
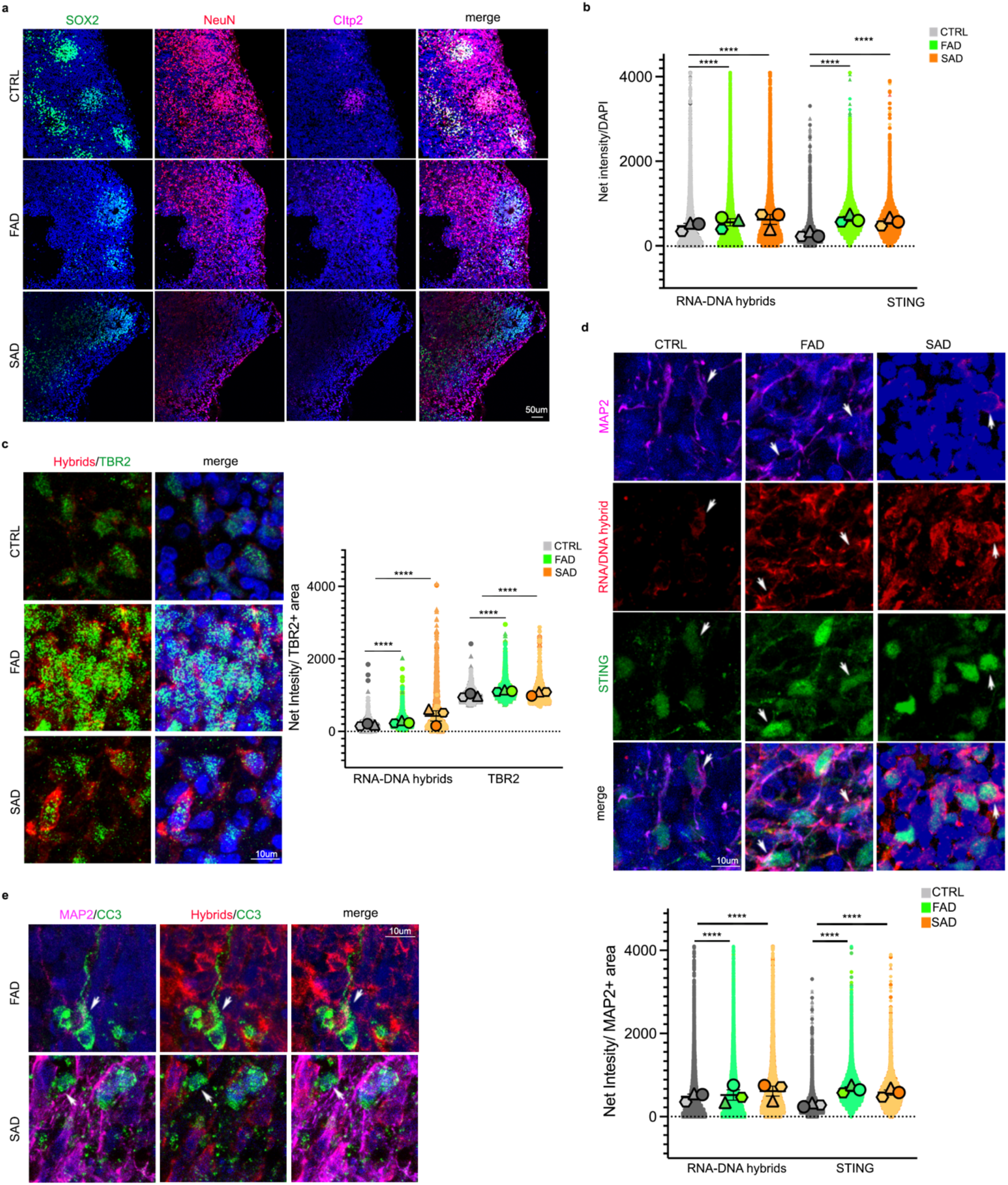
Characterization of TE-derived RNA-DNA hybrids inducing the cGAS-STING cell-death axis in AD cerebral organoids. **a**. Immunofluorescence for progenitors (SOX2, green signal), immature neurons (NeuN, red signal), and mature neurons (Citp2, pink signal) in CTRL and AD (FAD and SAD) organoids. Scale bar 50um, 20X magnification. DAPI staining on nuclei in blue. **b**. Quantification of the immunofluorescences in Figure 7b. **c**. Immunofluorescence for intermediate progenitors (TBR2, green signal) and RNA-DNA hybrids (red signal) in CTRL and AD (FAD and SAD) organoids. Scale bar 10um, 60X magnification 4X digital zoom. DAPI staining on nuclei in blue. **d**. Immunofluorescence for neurons (MAP2, pink signal) and RNA-DNA hybrids (red signal) and STING (green signal) in CTRL and AD (FAD and SAD) organoids. Scale bar 10um, 60X magnification 4X digital zoom. DAPI staining on nuclei in blue. **e**. Immunofluorescence for neurons (MAP2, pink signal), RNA-DNA hybrids (red signal) and cleaved caspase 3 (CC3; green signal) in FAD and SAD organoids. White arrows indicate MAP2/RNA-DNA hybrid/CC3-positive neurons. Scale bar 10um, 60X magnification 4X digital zoom. DAPI staining on nuclei in blue.

**Extended Data Table1.** TMPs values of genes involved in the WNT pathway that are differentially expressed in FAD hpNPCs.

|  | CTRL | FAD |
| --- | --- | --- |
| DKK1 | 3.692171 | 9.72627767 |
| DKK3 | 57.7179857 | 223.367699 |
| JUN | 77.3016 | 109.701667 |
| RAC2 | 6.636084 | 51.6482917 |
| RORA | 2.65834244 | 7.06845833 |
| SFRP4 | 0.8792335 | 14.3325863 |
| WNT2 | 0.01490205 | 3.81743227 |
| WNT5A | 59.2553505 | 204.736115 |
| WNT7A | 0.0380094 | 0.7898473 |

**Extended Data Table2.**
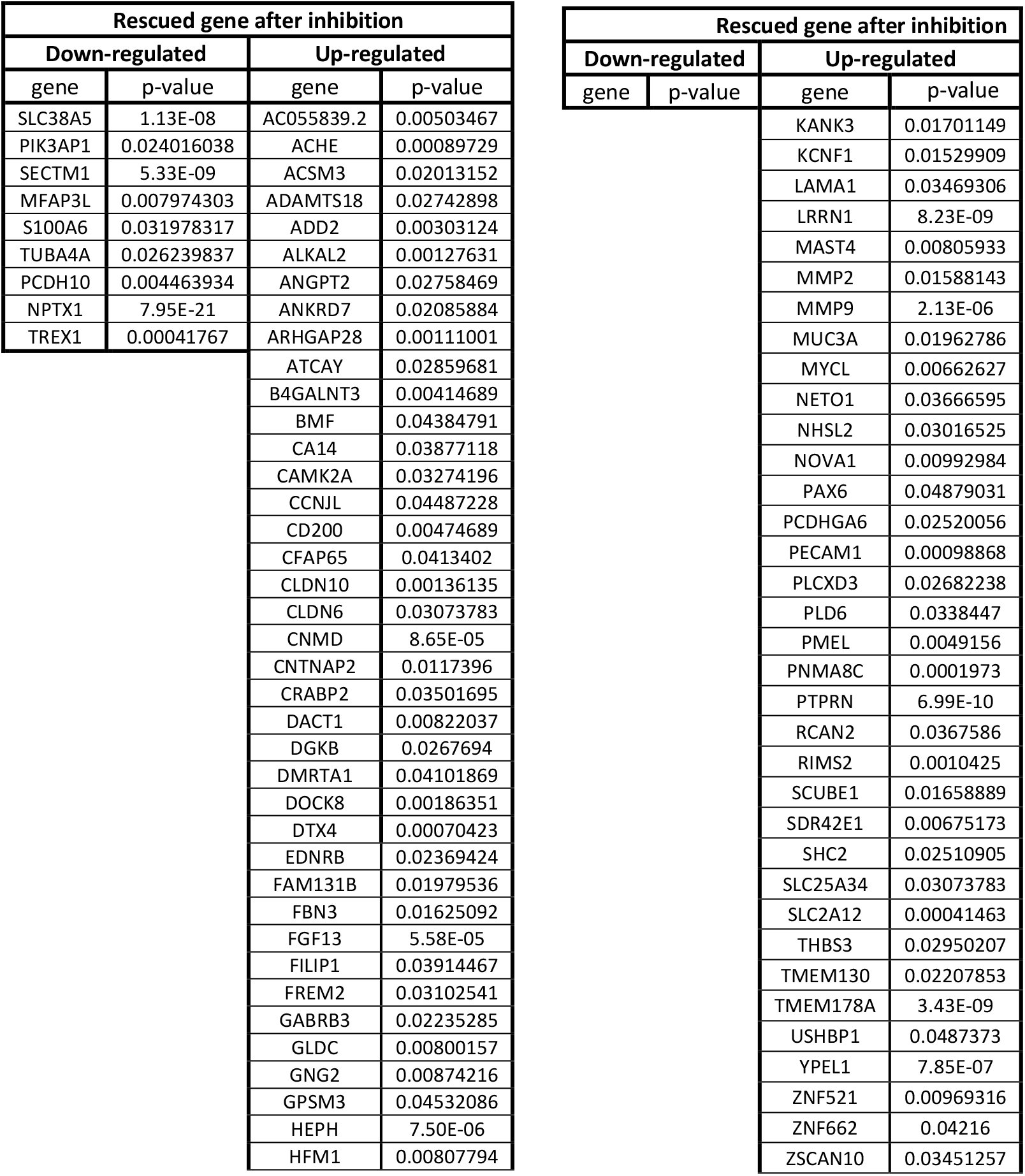
List of 83 completely rescued genes after the inhibition of c-JUN phosphorylation.

**Extended data Table 3.** Table of antibodies used.

| Antibody | Specificity | Host specie | Dilution | Supplier | Reference |
| --- | --- | --- | --- | --- | --- |
| OCT4 (MOUSE)<br>Clone 3A2A20<br>unconjugated | OCT4 | mouse | 1:200 | StemCell Technologies | 60093 |
| Human Nanog<br>Antibody | NANOG | goat | 1:20 | R&D Systems | AF1997 |
| c-Jun Monoclonal<br>Antibody (4H9) | cJUN | mouse | 1:1000 WB | Fisher | MA5-15889 |
| Anti-h nestin AF488<br>25 ug | NESTIN | mouse | 1:200 | Invitrogen | 5016830 |
| Tbr2 Abcam<br>antibody | TBR2 | rabbit | 1:200 | Abcam | ab216870 |
| Phospho-c-Jun<br>(Ser73) Polyclonal<br>Antibody | Phospho cJUN | rabbit | 1:1000 | Fisher | #44-292G |
| S9.6 | RNA-DNA hybrids | mouse | 1:500 | Kerafast | ENH001 |
| STING Polyclonal<br>Antibody | STING | rabbit | 1:100 IF<br>1:1000 WB | ThermoFisher | PA5-23381 |
| Cleaved Caspase-3<br>(Asp175) (5A1E)<br>Rabbit antibody | Cleaved caspase 3 | rabbit | 1:200 IF<br>1:1000 WB | CellSignaling | 9664T |
| SCGN Polyclonal<br>Antibody | SCGN | rabbit | 1:250 | Fisher | PA5-30393 |
| GRIK4 Monoclonal<br>Antibody (8H5G5) | GRIK4 | mouse | 1:100 | Fisher | MA5-31745 |
| Goat Polyclonal<br>Doublecortin<br>antibody | DCX | goat | 1:200 | Rockland Immunochemicals | 600-101-MH8 |
| AB5603 \| Anti-Sox2<br>(rabbit polyclonal) | SOX2 | rabbit | 1:1000 | EMD Millipore | AB5603-100UG |
| RBFOX3/NeuN<br>Antibody (1B7) | NeuN | mouse | 1:1000 | Novus Biologicals | NBP1-92693 |
| Mabe1045 \| Anti-CTIP2/BCL11B<br>Antibody, clone<br>25B6, Anti-CTIP2/BCL11B<br>Antibody, clone<br>25B6 | Cipt2 | rat | 1:1000 | EMD Millipore | MABE1045 |
| Anti-cGAS Antibody | cGAS | rabbit | 1:1000 | Millipore Sigma | ABF124 |
| MAP2 antibody | MAP2 | chicken | 1:1000 | Novus Biologicals | NB300-213 |
| GAPDH antibody<br>(rabbit) D16H11 | GADPH | rabbit | 1:1000 | Cell Signaling Technologies | 5174S |
| c-Jun (60A8) Rabbit<br>mAb | cJUN | rabbit | 1:200 IF | Cell Signaling Technologies | 9165 |
| Horse Anti-mouse<br>IgG, HRP-linked | mouse IgG | horse | 1:10000 | Cell Signaling Technologies | 7076S |
| Anti-rabbit IgG,<br>HRP-linked | rabbit IgG | goat | 1:10000 | Cell Signaling Technologies | 7074S |
| Cy™3 AffiniPure<br>Donkey Anti-Rabbit<br>IgG (H+L) | rabbit IgG | Donkey | 1:250 | Jackson ImmunoResearch | 711-165-152 |
| Alexa Fluor® 594<br>AffiniPure Donkey<br>Anti-Mouse IgG<br>(H+L) | mouse IgG | Donkey | 1:500 | Jackson ImmunoResearch | 715-585-150 |
| Alexa Fluor® 488 AffiniPure Donkey Anti-Rabbit IgG | rabbit IgG | Donkey | 1:500 | Jackson ImmunoResearch | 711-545-152 |
| Alexa Fluor® 488 AffiniPure Donkey Anti-Goat IgG (H+L) | Goat IgG | Donkey | 1:500 | Jackson ImmunoResearch | 705-545-003 |
| Alexa Fluor® 647 AffiniPure Donkey Anti-Mouse IgG (H+L) | mouse IgG | Donkey | 1:500 | Jackson ImmunoResearch | 715-605-150 |
| Goat anti-chicken IgY (H+T) Alexa Fluor Plus 647 | Chicken IgG | Goat | 1:1000 | Invitrogen | A32933 |
| Alexa Fluor® 647 AffiniPure Donkey Anti-Rat IgG (H+L) | Rat IgG | Donkey | 1:500 | Jackson ImmunoResearch | 712-605-150 |

**Extended Data Table 4.**
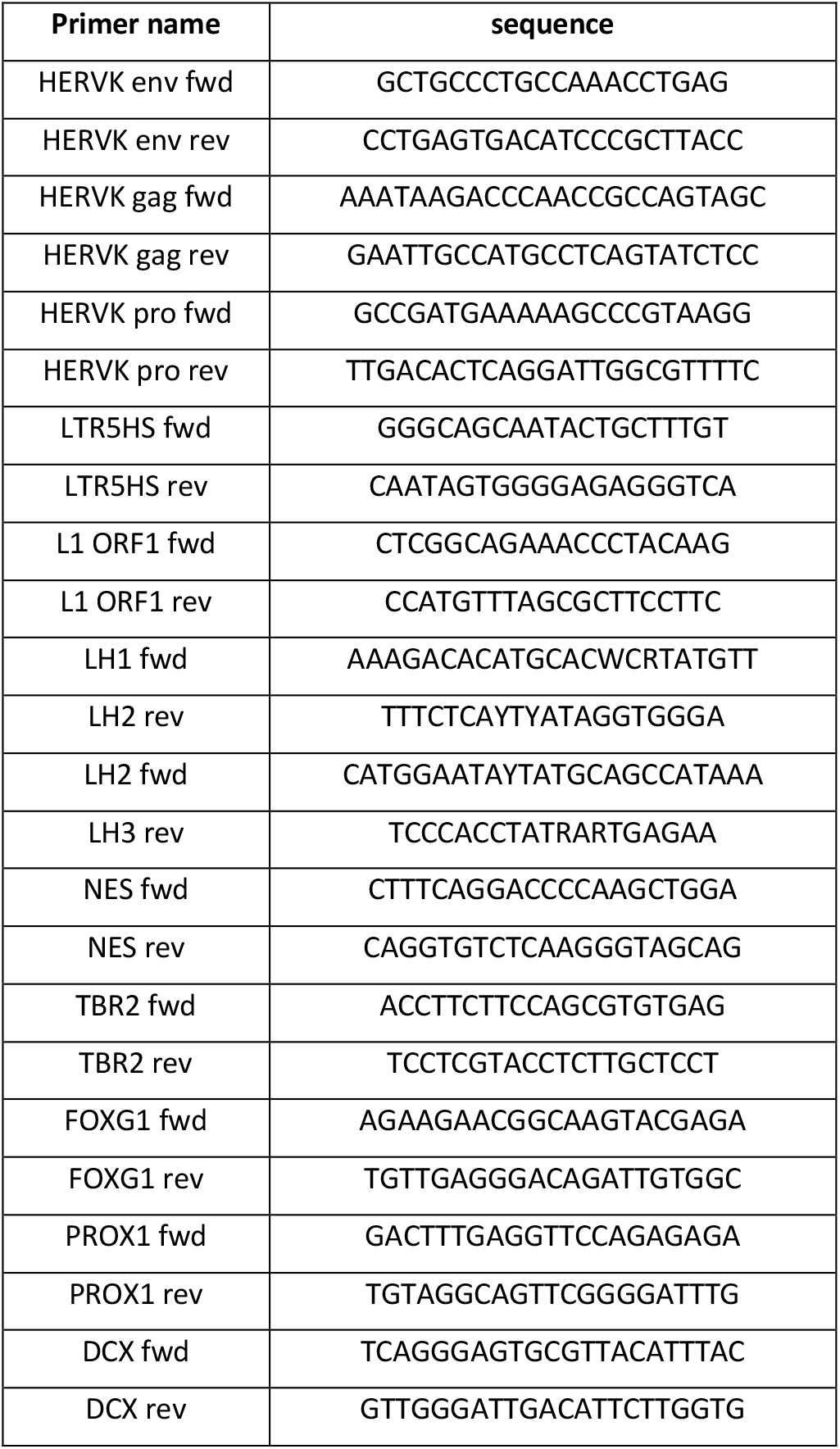
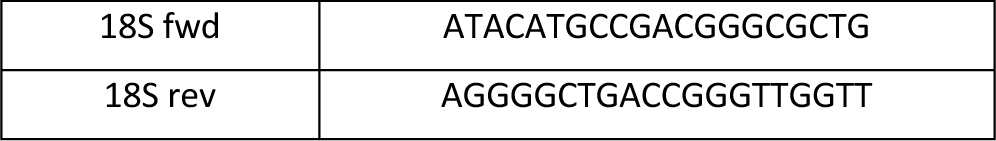
List of primers used for qRT-PCR analysis.

